# Natural variation in infection specificity of *Caenorhabditis briggsae* isolates by two RNA viruses

**DOI:** 10.1101/2024.02.10.579610

**Authors:** Cigdem Alkan, Gautier Brésard, Lise Frézal, Aurélien Richaud, Albane Ruaud, Gaotian Zhang, Marie-Anne Félix

## Abstract

Antagonistic relationships such as host-virus interactions potentially lead to rapid evolution and specificity in interactions. The Orsay virus is so far the only horizontal virus naturally infecting the nematode *C. elegans*. In contrast, several related RNA viruses infect its congener *C. briggsae*, including Santeuil (SANTV) and Le Blanc (LEBV) viruses. Here we focus on the host’s intraspecific variation in sensitivity to these two intestinal viruses. Many temperate-origin *C. briggsae* strains, including JU1264 and JU1498, are sensitive to both, while many tropical strains, such as AF16, are resistant to both. Interestingly, some *C. briggsae* strains exhibit a specific resistance, such as the HK104 strain, specifically resistant to LEBV. The viral sensitivity pattern matches the strains’ geographic and genomic relationships. The heavily infected strains mount a seemingly normal small RNA response that is insufficient to suppress viral infection, while the resistant strains show no small RNA response, suggesting an early block in viral entry or replication. We use a genetic approach from the host side to map genomic regions participating in viral resistance polymorphisms. Using Advanced Intercrossed Recombinant Inbred Lines (RILs) between virus-resistant AF16 and SANTV-sensitive HK104, we detect Quantitative Trait Loci (QTLs) on chromosomes IV and III. Building RILs between virus-sensitive JU1498 and LEBV-resistant HK104 followed by bulk segregant analysis, we identify a chromosome II QTL. In both cases, further introgressions of the regions confirmed the QTLs. This diversity provides an avenue for studying virus entry, replication, and exit mechanisms, as well as host-virus specificity and the host response to a specific virus infection.

## Introduction

Pathogens such as viruses exert strong selective pressures on their host. This may build into an ‘arms race’ of fast evolution between host immune defenses and viral counter-defense (Thompson 2005; Tenthorey et al. 2022). The rapid evolution of the virus is favored by a high mutation rate and large population size, and may lead to host specialization at the expense of host range (Barrett et al. 2009; Longdon et al. 2014; Rothenburg and Brennan 2020). Mechanistic studies of host range are well studied in plants (Barrett et al. 2009; Moffett 2009), as well as for viral host jumps to humans (Pepin et al. 2010; Woolhouse et al. 2012; Lu et al. 2015), but less using genetic approaches with a model animal. Here we address the host range variation for two related intestinal RNA viruses in the nematode species *Caenorhabditis briggsae*, a relative of *C. elegans* which can similarly be used for genetic studies.

The discovery of viruses naturally infecting *Caenorhabditis* species (Félix et al. 2011; Franz et al. 2012) provided an excellent genetic model to discover host factors and mechanisms of antiviral immunity. These natural *Caenorhabditis* viruses are bipartite, positive-strand RNA viruses related to the *Nodaviridae* family through both their RNA-dependent RNA polymerase (RdRP) and their capsid sequences. They infect intestinal cells and are transmitted horizontally through the fecal-oral route (Franz et al. 2014). Locally in France where the viruses were isolated, *C. elegans* and *C. briggsae* are the two dominant *Caenorhabditis* species found in rotting vegetal matter (Félix and Duveau 2012; Richaud et al. 2018; Frézal and Félix 2015). The Santeuil virus (SANTV) was the first virus to be discovered, in *C. briggsae*, through its effect on intestinal cells. The Orsay virus (ORV) was then found in *C. elegans* causing similar, but weaker, symptoms (Félix et al. 2011). A third virus was then found in *C. briggsae*, called Le Blanc virus (LEBV) (Figure 1A) (Franz et al. 2012). In nature, the Orsay virus (ORV) infects *C. elegans*, whereas Santeuil (SANTV) and Le Blanc (LEBV) viruses infect *C. briggsae* (Félix et al. 2011; Franz et al. 2012; Frézal et al. 2019). Despite this species-specificity, LEBV and SANTV are not particularly closely related compared to ORV (Frézal et al. 2019). A recently discovered fourth virus, the Melnik virus (MELV), was found in *C. briggsae* and is a close relative of SANTV (Frézal et al. 2019).

**Figure 1.**
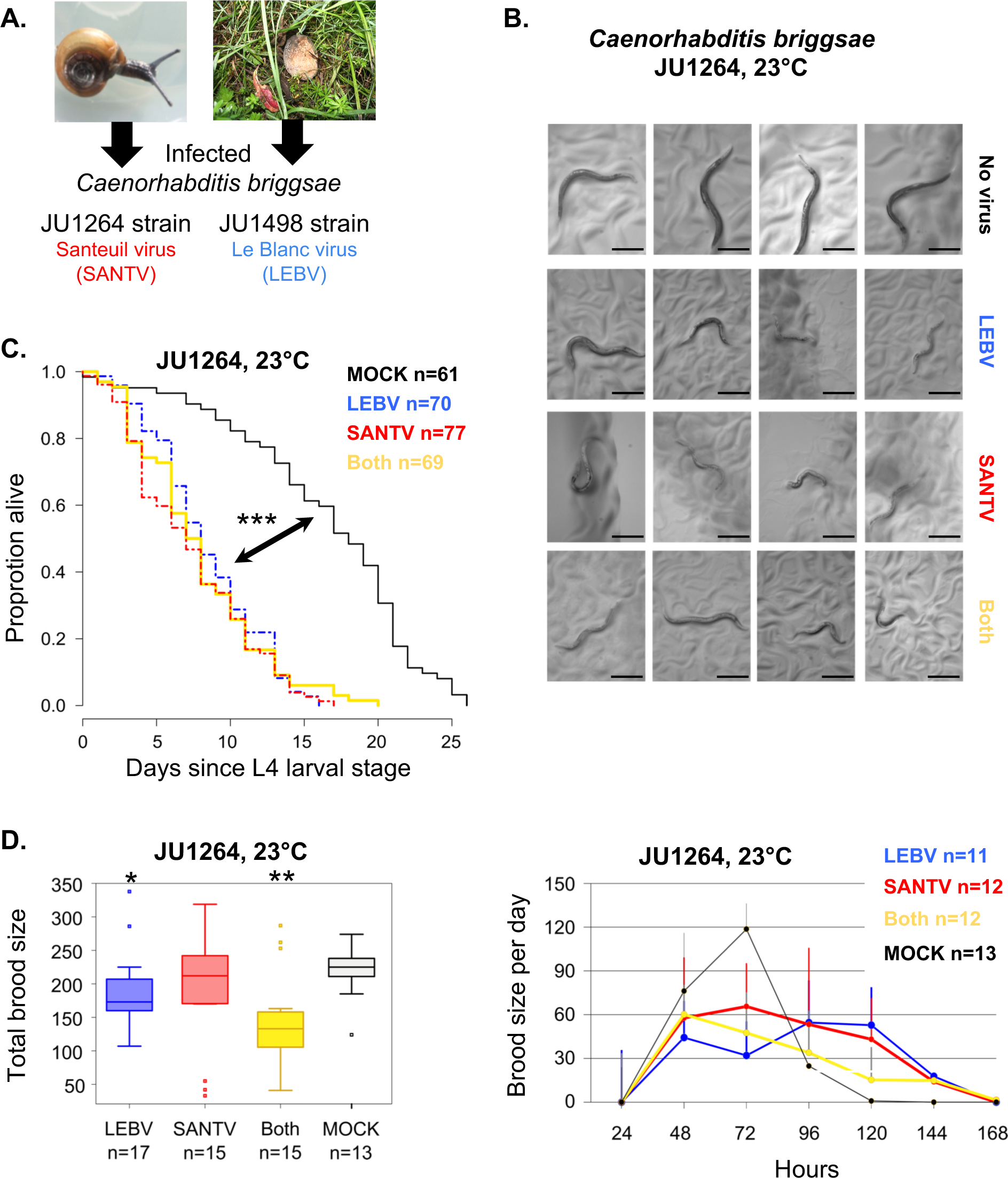
SANTV and LEBV delay and reduce *C. briggsae* progeny production in laboratory conditions. (A) The Santeuil virus (SANTV, strain JUv1264) was initially found in the *C. briggsae* JU1264 strain, isolated from a snail. The Le Blanc virus (LEBV, strain JUv1498) was found in *C. briggsae* JU1498 from a rotting peach (Félix et al. 2011; Franz et al. 2012). (B-E) The JU1264 strain was bleach-treated and infected by SANTV, LEBV or both. (B) Examples of infected animals showing a paler coloration of the gut compared to an uninfected *C. briggsae* adults and smaller body size or developmental delay at 120 hrs. Bars: 0.25 mm. (C) The viruses shorten the lifespan of *C. briggsae* JU1264 (logrank test, *p*=5.44 × 10^−9^, *p*=5.21 × 10^−10^, *p*=1.08 x 10^-7^, for mock *vs* LEBV, SANTV and both, respectively). (D) Total brood size (left) and number of embryos laid over time (right) in the same experiment. Compared to mock infection, the total brood size is smaller for LEBV infection and co-infection (Wilcoxon rank sum test with continuity correction, *p*=0.04 and *p*=0.01, respectively). JU1264 animals show a significant delay in progeny production when infected with either virus (linear model, *p*=2.2 × 10^−16^). ***: *p*<0.001,**: *p*<0.01,*: *p*<0.05.

*C. elegans* (and its congener *C. briggsae*) stands out as an invaluable organism for genetic investigations. They reproduce through selfing XX hermaphrodites and facultative X0 males for outcrossing. Given the model organism status of *C. elegans*, studies so far have focused on its interaction with ORV (Félix and Wang 2019). Natural collections of these species also allow for investigations of natural variation, using genome-wide association and recombinant genetic crosses between diverse wild strains to trace the genetic basis of phenotypes of interest (Andersen and Rockman 2022). Using natural variation to identify the genetic basis of host-pathogen interactions has two combined aims: discover molecular and cellular mechanisms at play, and analyze their evolutionary dynamics. The fast evolution of host-pathogen interactions makes this approach particularly powerful. Indeed, *C. elegans* wild isolates were found to strongly differ in their ability to replicate the ORV when infected in the laboratory. A genome-wide association study (GWAS) revealed an intermediate-frequency indel polymorphism in the *drh-1* gene (encoding a RIG-I-like helicase) as the main locus explaining variation in ORV sensitivity (Ashe et al. 2013). This RIG-I homolog triggers viral genome degradation via small RNA silencing pathways (Ashe et al. 2013; Guo et al. 2013) and a host transcriptional response (Sarkies et al. 2013; Bakowski et al. 2014; Chen et al. 2017; Tanguy et al. 2017; Sowa et al. 2020; Lazetic et al. 2022). In addition to this genome-wide association approach, a biparental cross between the reference strain N2 and the wild isolate CB4856 recently identified a quantitative trait locus on the right of chromosome IV, which may be partially explained by a non-synonymous polymorphism in the *cul-6* gene, coding for a cullin, an ubiquitin ligase cofactor (Sterken et al. 2021).

In addition to the analysis of natural variation, forward and reverse genetic studies in *C. elegans* revealed a number of factors required in host defense or for the viral cycle (Jiang et al. 2017; Tanguy et al. 2017; Le Pen et al. 2018; Long et al. 2018; Casorla-Perez et al. 2022; Cubillas et al. 2023). Especially, *C. elegans* antiviral immune response involves: i) the small RNA response; ii) the ubiquitin pathway (Bakowski et al. 2014; Chen et al. 2017); iii) a conserved SID-3-dependent signaling pathway involved in receptor-mediated endocytosis, similar to their mammalian orthologs (Jiang et al. 2017), and in the phosphorylation of STA-1, a homolog of mammalian STAT (Tanguy et al. 2017); iv) the conserved role of uridylation in destabilization of the viral RNA (Le Pen et al. 2018).

ORV was recently shown to enter and initially replicate in other *Caenorhabditis* species, but not *C. briggsae* (Shaw and Kennedy 2022). By contrast, SANTV and LEBV both undergo full viral cycles in *C. briggsae*. They thus provide a model to study the pattern of sensitivity within the host species for two different viruses and hence of viral specificity, which is the focus of the present work.

Infecting a panel of 40 *C. briggsae* natural isolates with SANTV and LEBV, we found that: 1) most *C. briggsae* isolates of tropical origin are resistant to both viruses (e.g. the reference strain AF16); 2) most isolates of temperate origin are sensitive to both viruses (e.g. the JU1264 or JU1498 isolates in which the original SANTV and LEBV were discovered); and 3) some are specifically sensitive to one virus (e.g. HK104 from Japan is specifically sensitive to SANTV and resistant to LEBV). We analyzed the small RNA response to each viral infection in JU1264 and HK104 and found that the sensitivity to SANTV and LEBV does not correspond to a defect in the small RNA response and amplification. Instead the difference between resistant and sensitive hosts appears to originate from variation at earlier steps of entry or replication of the viruses. We then focused on the HK104 strain showing viral infection specificity and investigated the quantitative genetic underpinnings of viral sensitivity using two panels of recombinant inbred lines built from pairwise crosses of *C. briggsae* isolates: 1) virus-resistant AF16 and SANTV-sensitive HK104; 2) virus-sensitive JU1498 and LEBV-resistant HK104. In the first case, we found two QTLs on chromosomes IV and III, and in the second a major QTL at the right end of chromosome II. This study grounds *C. briggsae* and its viruses as an interesting pathosystem to study the sensitivity and specificity of host-viral interactions.

## Materials and methods

### Culture and strains

A list of *C. briggsae* and *C. elegans* strains used in this study can be found in Table S1. *C. briggsae* was grown under standard conditions used for *C. elegans* (Stiernagle 2006), but at 23°C unless otherwise indicated. A bleach treatment was applied to all isolates prior to virus infection to eliminate possible contaminations, as in Stiernagle (2006) and Félix et al. (2011). Virus filtrates were prepared as described previously (Félix et al. 2011). The viral isolates JUv1498 was used for LEBV and JUv1264 for SANTV, except in the small RNA experiment where SANTV JUv1993 (Frézal et al. 2019) was also used.

### Longevity and brood size assays

For the assays in this section, we used Normal Growth Medium with a higher agar concentration to prevent animals from burrowing (NGM with 2.5% agar). Plates were seeded with *E. coli* OP50 (Stiernagle 2006).

Initially infected JU1264 *C. briggsae* hermaphrodite cultures were generated by plating 5 L4 stage larvae from an uninfected culture onto fresh plates and adding 50 μL of either sterile ddH_2_O (mock), a filtrate of SANTV, LEBV or a mixture of 50-SANTV:50-LEBV filtrates to the plates. Each inoculated culture was maintained for 4 days at 23°C by transferring a 0.5 cm^2^ square of agar to 3 new plates with food on day 3 and then day 5. The success of infection was checked by FISH for all plates of infected conditions.

For each of the four conditions (mock, SANTV, LEBV, SANTV-&-LEBV infections), the longevity assay was started by transferring 80 L4 stage larvae from the inoculated cultures onto 4 plates (20 animals per plate; 2 or 3 from each of the nine starting cultures). Each pool of 20 adult animals was carefully transferred to a new plate every day until the end of the reproductive period. Death was recorded when the animal did not react to prodding with a worm pick.

For the brood size assay, 20 L4 animals were isolated for each treatment (one animal per plate, 2 or 3 randomly picked from the nine starting cultures). Each animal was then transferred to a new plate every 24 hrs. Each scored individual was transferred to a new plate 24, 48, 72, 96,120, 144 and 168 hrs after the L4 stage. Progeny numbers were scored 48 hrs after each transfer. To ease scoring, some plates were cooled to 4°C after 48 hrs and scored within two days. Note that *C. briggsae* animals often disappear by burrowing into the NGM agar, which explains that replicates are missing from the data when compared to the initial number of animals.

### Infection of the set of *C. briggsae* natural isolates

We performed three separate tests (Batches 1-3 in Figure S2A) of infection of the same set of *C. briggsae* strains, and in each batch the infection of each virus was performed in duplicate or triplicate (Figure S2). Before viral inoculation, 10 L4 stage larvae of a previously bleached culture were placed onto 55 mm NGM plates seeded with *E. coli* OP50 (Stiernagle 2006). 30-40 μL of filtrates of the viruses were added into the middle of the *E. coli* OP50 lawn. Inoculated cultures were incubated at 23°C for 7-8 days. Maintenance of the infected cultures was performed by transferring a piece of agar every 2-3 days to a new plate with food. At 7-8 days post-infection, nematodes from two plates were collected in Ultrapure water (Invitrogen) for the FISH assay (see below).

### Fluorescent in situ hybridization (FISH) for viral RNA

The protocol is adapted from Raj et al. (2008) as in Franz et al. (2014) with the multiple probes per viral RNA molecule used in Frézal et al. (2019). Infected nematodes were harvested with Nanopure nuclease-free water (Invitrogen) and pelleted by centrifugation for 5 min at 3000 rpm in 15 mL Falcon tubes. The pellet was transferred to a non-adhesive 1.5 mL tube (Axygen). 1 mL of fixative solution - consisting of the following: 5 mL 37% formaldehyde (Sigma #533998), 5 mL 10x PBS (Ambion AM962, pH 7.4), and 40 mL nuclease-free water (Invitrogen) - was added and rotated at room temperature with light agitation for 40 min. After removing the fixative solution, the pellet was washed twice with 1x PBS (Invitrogen), resuspended in 70% ethanol and stored at 4°C. After at least overnight at 4°C, the nematodes were pelleted and washed with the wash solution consisting of the following: 10 mL deionized formamide (Ambion AM9342), 5 mL 20X SSC (Ambion AM9770) and 35 mL nuclease-free water (Invitrogen). Nematodes were then suspended in 100 μl of the hybridization buffer consisting of the following: for 10 mL, 1 g dextran sulfate (Sigma #D6001), 1 mL 20x SSC, 2 mL deionized formamide, and 7 mL nuclease-free water (Invitrogen). 1 µL of 1:40 diluted fluorescent probes were added and the nematodes incubated overnight at 30°C in the dark. The next day, they were washed with 1 mL wash solution, resuspended in 0.02% DAPI (Sigma #D9564, 5 mg/mL) in 1 mL wash solution and incubated 30 min at 30°C in the dark. Finally, they were suspended in 2x SSC and kept at 4°C for imaging. For imaging, ∼4 µL of sample were pipetted onto round coverslips, then sealed onto glass slides using silicon isolators. The fluorescence was visualized directly with an Olympus FV1000 macroscope (Batches 1 and 2) or with a Zeiss AxioImager M1 (Batch 3). Only adults were scored.

SANTV infected animals were scored using the RNA1 probes labeled with Alexa (Cal fluor Red 610) (Frézal et al. 2019). We used Alexa RNA2 probes for LEBV in experimental Batches 1 and 2 and Cy5 (Quasar 670) RNA1 probes for LEBV in Batch 3 (Frézal et al. 2019). Representative images are shown in Figure S2B; a grand mean of results, provided in Table S4, was calculated for Figure 2B. The proportion of infected animals were overall higher in Batch 3 but the qualitative results are similar.

**Figure 2.**
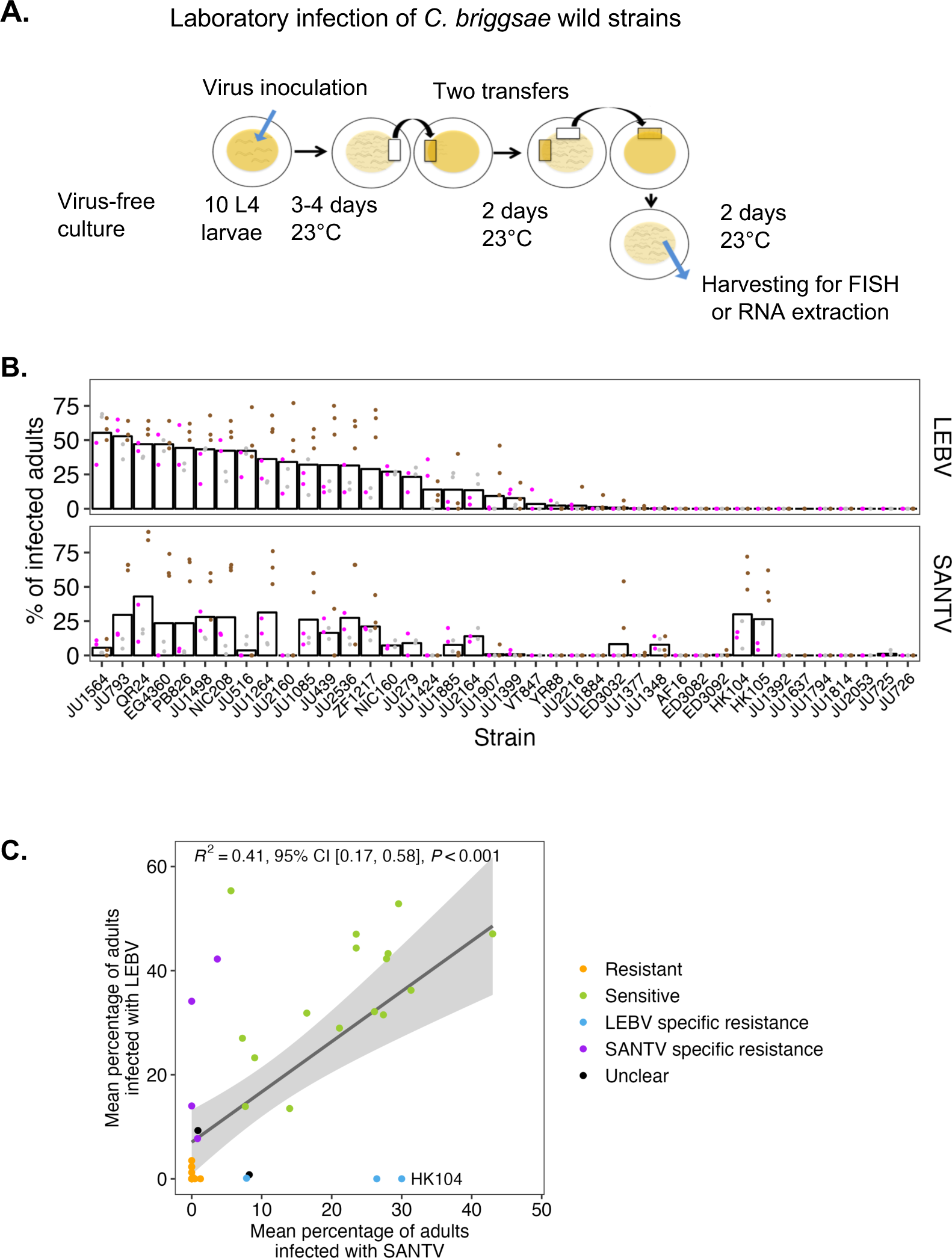
Variation in sensitivity to SANTV and LEBV.of *C. briggsae* wild isolates. (A) Experimental infection protocol, starting from a bleached *C. briggsae* culture. (B) Sensitivity of *C. briggsae* wild isolates to SANTV and LEBV. The two viruses were inoculated in parallel. This graph represents the percentage of infected hosts as assayed by FISH for the corresponding virus. Dots are replicates within a block, with 100 animals scored per replicate (see Table S4 for the detailed results and Figure S2 and Methods for the experimental design). Experimental blocks are represented by colors and the bar indicates the grand mean of the blocks. The strains on the *x* axis are ordered by their rank of LEBV sensitivity. (C) Two-dimensional plot displaying the grand mean of panel B for SANTV and LEBV. The dots represent individual strains that are colored by categories. Their sensitivity to each virus is coded as a binary trait and the combination color-coded. The variance among replicates was considered, which explains that strains with similar positions on this plot are differently colored, one being labeled “Unclear”. The strains show a significant correlation between their sensitivity to SANTV and LEBV (regression line in dark grey, with the 95% confidence interval in light grey). In addition, this plot highlights the specificity of infection for some strains, such as HK104, located far outside this diagonal..

### Haplotype network of *C. briggsae* wild isolates

For the network in Figure 3B, we obtained whole-genome sequence data of 39 wild *C. briggsae* strains from *Caenorhabditis* Natural Diversity Resource (CaeNDR) (Crombie et al. 2023) and called genetic variants among them using the pipeline *wi-gatk* (https://github.com/AndersenLab/wi-gatk). We pruned the resulting hard-filtered VCF to 1,958,505 biallelic SNVs without missing genotypes using *BCFtools* (v.1.9) (Li 2011). Then, we converted this pruned VCF file to a PHYLIP file using the *vcf2phylip.py* script (Ortiz 2019). The haplotype network was built from 1,958,505 informative sites using SplitsTree4 (Huson & Bryant, 2006).

**Figure 3.**
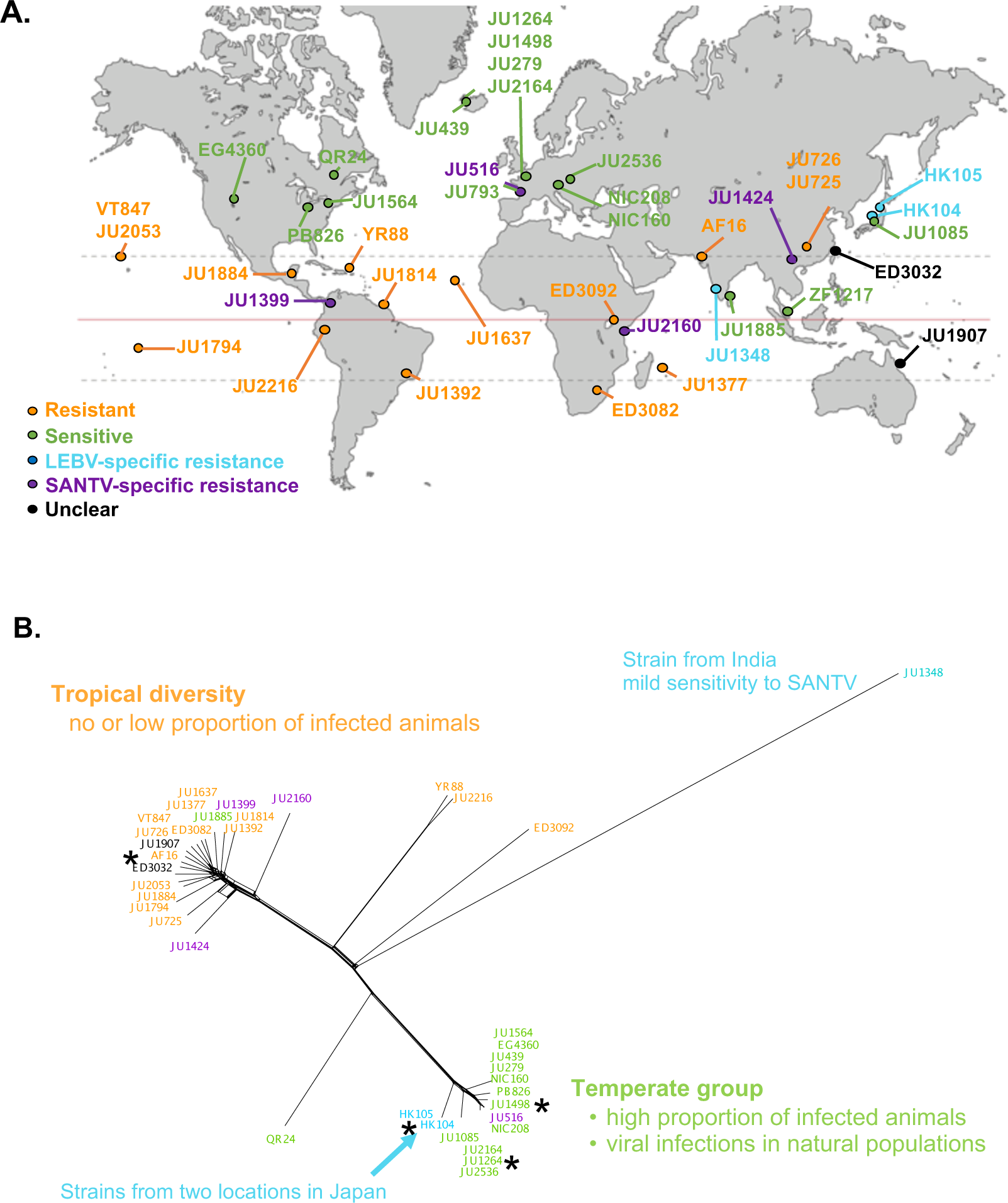
Relationship between *C. briggsae* virus sensitivity, geographic origin and genetic relatedness. (A) World map showing the geographical distribution of the *C. briggsae* strains tested in Figure 2. Their sensitivity to both viruses is color-coded as in Figure 2C. In determining specific resistance, a low level of viral replication was considered as indicating some sensitivity, thus neglecting for the sake of simplicity possible quantitative variation in sensitivity between the two viruses. For example, JU1564 and JU1907 could be included as specifically resistant to SANTV. (B) Genetic relationship represented by a haplotype network between the *C. briggsae* isolates, based on available genome sequences at CaeNDR. * indicate strains used in further studies.

### RT-qPCR for viral RNA

RT-qPCR was performed to analyze viral load in the *C. briggsae* populations in addition to the FISH assay which measures a proportion of infected animals. The mixed-stage nematode population from two Petri dishes was pelleted in M9 solution and frozen at −20°C. RNAs were extracted by adding 50 μL Trizol and vortexing. The mix was frozen in liquid nitrogen, then transferred to a 37°C waterbath; this freeze-thaw procedure was repeated four times. The tube was vortexed at room temperature for 30 sec then left to settle for 30 sec; this vortexing procedure was repeated five times. 10 μL chloroform were then added, mixed by manual agitation of the tube for 15 sec and incubated at room temperature for 3 min. The tubes were centrifuged at ca. 10,000 g for 15 min at 4°C. The aqueous phase containing RNAs was transferred to a new tube and 25 μL isopropanol were added, followed by a 10-min incubation at room temperature. The mix was centrifuged and the supernatant discarded. The pellet was washed with 75% ethanol, vortexed and centrifuged (7,000 g, 5 min, 4°C). The supernatant was discarded and the pellet let to dry with opened lid. The nucleic acids were dissolved with 20 μL RNAse free ddH_2_O, incubated in a 55°C waterbath for 3 min to help resuspension and stored at −80°C.

This nucleic acid preparation was then thawed and treated with DNAse (Invitrogen) at 37°C for 20 min in a final volume of 5.8 μL, after which 0.2 μL EDTA at 0.25 M was added to stop the reaction (75°C for 10 min).

cDNA was generated from total RNA with specific primers using Superscript III (Life Technologies). For the RT-PCR experiments (not quantitative), the annealing temperature was 64°C. For the RT-qPCR experiments, the SYBR Green I Master mix was used on a LightCycler 480 Real Time PCR System (Roche). The primers are listed in Table S2. The results were normalized to *Cbr-eft-2* expression in the same sample and then to the viral RNA level in another sample of infected JU1264 (SANTV) or JU1498 (LEBV) animals.

### Preparation of small RNA libraries

200 embryos of each strain (AF16, HK104 and JU1264) were suspended in nuclease-free water and plated onto 55 mm NGM plates seeded with *E. coli* OP50. Once the liquid was absorbed by the agar, 50 μL of SANTV or LEBV filtrate, or both filtrates were added to the plates. After inoculation, plates were incubated for 3 days at 23°C. Adult hermaphrodites were then harvested with M9 (Stiernagle 2006) and washed twice in UltraPure water (Invitrogen). After the animals had pelleted under gravity, 800 μL of TRIzol (Invitrogen) were added to each worm pellet and the mixes were snap-frozen in liquid nitrogen before being stored at −80 °C. The next day, total RNAs were extracted by adding 200 μL of chloroform to the mix. After a 15-minute centrifugation step at 13,000 rpm, the upper phase was collected. This step was repeated twice. RNAs were then precipitated by the addition of 500 μL isopropanol and 1 μL glycogen and an overnight incubation at −20°C. The next day RNAs were pelleted at 13,000 rpm for 30 min and the pellets washed twice in 75% ethanol. The pellets were air dried and dissolved in nuclease-free water. RNA concentrations were quantified using the Nanodrop (Thermofisher)

To generate 5’-end independent small RNA libraries, 800 ng of total RNAs were treated with 5’-polyphosphatase (Epicenter/Illumina) for 30 min. Libraries were generated using the NEBNext® Small RNA Library Prep Set for Illumina® following the manufacturer’s instructions from step 1 to 15. Migration on denaturing polyacrylamide (Novex™ TBE-Urea Gels 6% from ThermoFisher) was used for the library size selection. The part of the gel located between the ladder fragments at 147 and 160 bp was extracted. Libraries QC was performed using Bioanalyser with the Agilent High Sensitivity DNA Kit before being sequenced using an Illumina NextSeq System machine to generate 75-nucleotide single-end reads. Reads are available at NCBI with accession number PRJNA1046456.

### Small RNA content analysis

The program Cutadapt v1 was used to remove adapters from the Fastq files to recover reads of length between 16 and 33 nucleotides. Reads were aligned onto viral genomes SANTV (JUv1264; NC_015069.1, NC_015070.1) and LEBV (JUv1498; NC_028134.1, NC_028133.1) and onto host genomes (*C. briggsae* WS238.genomic_masked.fa; *C. elegans* WS245.genomic.fa available at: https://downloads.wormbase.org/species/c_elegans/sequence/genomic/ and https://downloads.wormbase.org/species/c_briggsae/sequence/genomic/) (Table S5). Counts of reads grouped according to their length, sense, and the identity of their first nucleotide were obtained with a shell script (Table S5, second sheet). Data were normalized to the total number of reads with a length between 16 to 33 nucleotides. Plots of the proportions of siRNA grouped according to their length, sense and their first nucleotide were generated to investigate the antiviral response of each *C. briggsae* host to the infection with LEBV, SANTV or both. Sequences are available under the NCBI project <u>PRJNA1046456</u> (SRR27205657-SRR27205668).

### Infection and QTL mapping using the Advanced Intercross Recombinant Inbred Lines (AI-RILs) between AF16 and HK104

We phenotyped 65 RILs between AF16 and HK104 (Ross et al. 2011) for their sensitivity to the Santeuil virus (Table S6). For each RIL, we infected 55-mm plates seeded with *E. coli* OP50 containing 10 L4 larvae, in triplicate. Cultures were incubated with 30 μL of SANTV JUv1264 filtrate at 23°C for 7 days as above. FISH was performed as above.

Phenotype data were coded as binary (infected or non-infected) or as a quantitative trait (percentage of infected animals). The R/QTL package was used for interval mapping with the following options: crosstype = “riself” and model = “binary” or “2part” (Broman et al. 2003), using the genotype data of the individual AI-RILs from Ross et al. (2011) integrated in the Cb4 assembly of *C. briggsae* AF16 (Ross et al. 2011; Davis et al. 2022). A two-qtl procedure was also employed with the “scantwo” option. The significance threshold LOD score was estimated via 10,000 permutation tests.

### Near Isogenic Lines (NILs) with AF16 and HK104

The NILs JU2831, JU2832, and JU2833 were constructed by backcrossing 3 AIRILs with the QTL region of either AF16 or HK104 into the background of the other parental strain (Figure 6, Table S10). The genotype at chromosome IV was followed using the SNP marker *cb13587* (Table S6). JU2915 and JU2916 were created by backcrossing the NIL JU2832 to AF16 to separate the QTL regions on chromosomes III and IV. NILs were genotyped using markers shown in Tables S2 and S10. Genotyping was performed by pyrosequencing using a PyroMark Q96 ID instrument from Biotage, according to the manufacturer’s instructions, as in Aydin et al. (2006) and Ashe et al. (2013). NILs were phenotyped for SANTV susceptibility following the same method as above.

### Construction and assay of Recombinant Inbred lines (RILs) between JU1498 and HK104

Reciprocal genetic crosses were performed between JU1498 and HK104 using hermaphrodite L4 larvae and males. Heterozygous larvae (stage L4) of the F1 cross progeny were singled onto 55-mm plates and allowed to self for 10 generations by randomly picking a single hermaphrodite each generation.

We phenotyped the RILs for their susceptibility to Le Blanc virus. For each RIL, we infected 55-mm plates seeded with *E. coli* OP50 containing 10 L4 larvae, in triplicate (Table S8). Cultures were incubated with 30 μl of Le Blanc virus filtrate at 23°C for 7 days. Maintenance of the infected cultures over more than 4 days was performed by transferring every 2–3 days a piece of agar to a new plate with *E. coli* OP50. At 7 days post-infection, nematodes from two plates were collected in Nanopure water (Invitrogen) for the FISH assay.

### Pool sequencing and QTL analysis with the RILs between JU1498 and HK104

From the 79 RILs, 37 resistant lines and 23 highly sensitive lines were chosen to represent the two extremes of the phenotypic distribution. Genomic DNA was extracted from mixed stage growing populations using reagents from the Puregene Core Kit A (QIAGEN, Valencia, CA) and quantified using Qubit 3.0 fluorometer (Life Technologies) with the dsDNA BR Assay Kit according to the manufacturer’s protocols. The sequencing was performed by BGI genomics with paired-end sequencing using IlluminaHiSeq 4000 at 30x coverage. Reads are available at NCBI with accession number PRJNA1053628.

Adapter sequences and low-quality reads in raw sequencing data of the two parents (HK104 and JU1498) and the two pools of RILs were removed using *fastp* (v0.20.0) (Chen et al. 2018). Then, we aligned the trimmed FASTQ files to the six chromosomes (I, II, III, IV, V, X) of the *C. briggsae* AF16 reference genome (WS283) (Davis et al. 2022) using *BWA* (Li and Durbin 2009) incorporated in the pipeline *alignment-nf* (https://github.com/AndersenLab/alignment-nf) (Cook et al. 2017). Single nucleotide variant (SNVs) were called for each generated BAM against the reference using *GATK* (v4.1.4) (Poplin et al. 2018) incorporated in the pipeline *wi-gatk* (https://github.com/AndersenLab/wi-gatk/) (Cook et al. 2017). We selected 14,749 SNVs in the hard-filtered VCF file by requiring full information of the four samples, different homozygous alleles between the parents, different alleles between the two pools, and numbers of mapped reads between 10 and 50 at each SNV in the two pools.

We performed bulk segregant analysis to detect genomic regions where parental allele proportions deviate between the two extremes, here between the Le Blanc virus sensitive and resistant pools as explained (Frézal et al., 2018). For this, we calculated the frequency of the HK104 allele at each SNV for each pool (Table S9). We further performed a sliding window analysis with a 50-SNVs window size and a one-SNV step size for the frequency of each pool.

To test whether differences in HK104 allele frequencies between the sensitive and resistant pools were significantly different from expectations under a random distribution, we first calculated for all previously defined windows the log-odds ratio as: log(m_1_/(n_1_-m_1_))/ (m_2_/(n_2_-m_2_))), m_1_ being the HK104 allele proportion multiplied by the number of RILs in the sensitive pool (n_1_=23) and m_2_ the HK104 allele proportion multiplied by the number of RILs in the resistant pool (n_2_=37). We calculated the threshold of significance (*p* = 0.01) in a two-tailed manner by simulating log-odds ratios for one million randomized draws of the two pools using the binomial law, as in Frézal et al. (2018).

To find molecular markers for genotyping, the Pindel software (Ye et al., 2009) was used to detect homozygous indels in the JU1498 and HK104 strains. Parent-specific deletions were identified and manually checked using Tablet (Milne et al., 2013).

### Near Isogenic Lines (NILs) between HK104 and JU1498

The NILs JU3241, JU3244, JU3245, JU3246, and JU3247 were first built in two cross directions to confirm the QTL region detected through the bulk segregant analysis. We crossed JU1498 L4 hermaphrodites with HK104 males. The male F1 cross progeny were crossed to either JU1498 or HK104 hermaphrodites, to introduce chromosome II of HK104 and JU1498, respectively, in the other background

To generate further recombinants, we crossed JU3247 hermaphrodites to HK104 males and screened with three pairs of primers: CbrII14,973F/ CbrII14,973R, CbrII15,198F/CbrII15,198R and CbrII16,567F/CbrII16,567R (Table S2). The recombinants obtained JU4033 and JU4034 were then genotyped using primers II:16,058-F/II:16,058-R (PCR for indel) and II:16146276-F/II:16146276-R (pyrosequencing).

Chromosome II ends at 16.627 Mb in WormBase Cb4 assembly of AF16 and at 16.595 Mb in QX1410 at CaeNDR. Both long-read assemblies (Ren et al. 2018; Stevens et al. 2022) introduce in the QTL region some genes, such as *Cbr-eri-3* located on another contig in Cb4, or *T23F4.2* that was placed further left. The QTL left bound at 16.05 Mb corresponding to 15.985 Mb on the Stevens et al. assembly.

### CRISPR-Cas9 genome editing of *CBG01824 and Cbr-rsd-2*

The Cas9 protein (IDT) was injected with RNA oligonucleotides including guide RNAs targeting the genomic region of interest. The CRISPR guide RNAs were designed using http://crispor.tefor.net/ and ordered from IDT.

*CBG01824*: The repair template for targeted replacement in the *CBG01824* gene was designed to replace the T nucleotide at position 10349333 in AF16 (Cb4) to a C as in HK104, which results in a Valine to Alanine amino acid change. To prevent a second cut after repair, a C to A synonymous mutation at position 10349346 was introduced to modify the NGG site. Lastly, to screen for the replacement, an A to T synonymous mutation at position 10349340 and a C to T synonymous mutation at position 10349343 were introduced so as to yield a different band pattern after restriction enzyme cutting. The list of RNA oligonucleotides and PCR primers are in Table S2.

Before injection, 1.5 μL of 200 μM guide RNA (synthesized by IDT) and 1.5 μL of 200 μM tracer RNA (IDT) were incubated in a PCR machine at 95°C for 3 minutes followed by decreasing temperature steps of 5°C steps every minute until 25°C. We used a guide RNA targeting the *Cbr-dpy-1* gene on chromosome III as a co-CRISPR marker (Culp et al. 2015). 1.0 μL of 100 μM *dpy-1* guide RNA and 1.0 μL of 100 μM tracer RNA were pre-incubated using the same temperature protocol. The final injection mix was: 3 μL of crRNA-tcRNA, 2 μL of dpy-1crRNA-tcRNA, 2.9 μl Cas9 (10 ng/μL), 2.5 μL of repair template (Eurofins), 0.44 μL of nuclease free water (Invitrogen), and 0.36 μl HEPES (IDT). This mix was pre-incubated at 37°C for 30 minutes and used within 3-5 hours.

Young adults (preferentially with 1-2 embryos in the uterus) were injected using the Eppendorf Transjector 5246. The injected animals were maintained at 25°C for 3-4 hours, transfered to new *E. coli* OP50 plates and kept at 25°C. If dumpy progeny (co-CRISPR marker) was seen on the P0 plates, the plate was selected and F1 progeny was singled. Once F2 progeny were seen on the F1 isolation plates, mixed-stage populations were pooled to perform a PCR targeting the region around the edit, using the GoTaq Master Mix (Promega) according to the manufacturer’s protocols. The PCR product was digested with a restriction enzyme using either FastDigest Bsp1407I or Pael (SphI) (Thermo Fisher) following the manufacturer’s protocols. The products were run on a 3-4% agarose gel at 60 V for 1 hour. Once the samples with the desired cut were detected, 8 or more worms were singled from the corresponding plate to select for homozygous gene edits. The PCR products with a desired edit were sent to Eurofins for Sanger sequencing for confirmation of the replacement.

A knock-out of the gene *CBG01824* in the AF16 background (JU3436) was also created using the same CRISPR protocol but without repair template, creating a small deletion. We also generated two *Cbr-rsd-2* knock-out mutants in exon 3 and 18 (JU3656 and JU4131) in JU3414 background to inactivate one and all putative isoforms, respectively, using the *C. elegans* gene annotation. The list of RNA oligonucleotides and PCR primers used for the screening of small indels are in Table S2. All edits were verified by Sanger sequencing.

## Results

### LEBV and SANTV infections delay *C. briggsae* progeny production

We first tested the effect of SANTV and LEBV infection and their co-infection on longevity and progeny production, in *C. briggsae* JU1264, the strain in which SANTV was initially isolated. We inoculated with either SANTV or LEBV, or both viruses, JU1264 animals from a culture that had first been bleached and then cultured on *E. coli* OP50. The infection status was monitored by the pale intestinal coloration of individuals (Figure 1B) and using fluorescent in situ hybridization (FISH) with which co-infected animals were visible, as shown before (Frézal et al. 2019). Infections by either virus shortened the host’s lifespan, although most animals still survived through the reproductive period (Figure 1C, Table S3). As before (Félix et al. 2011), we did not see a significant effect of SANTV infection on the total brood size, but LEBV and especially the co-infection significantly lowered brood size (Figure 1D). In exponentially growing populations, a reproductive delay may strongly decrease fitness: a key fitness consequence was the slowing down of progeny production, as previously shown for SANTV.

Isolates of LEBV were solely found in *C. briggsae* natural populations so far (Frézal et al. 2019). To test whether LEBV infections could be sustained in *C. elegans*, we inoculated a set of *C. elegans* isolates with LEBV. We could not detect any infection after 7 days using RT-PCR (Figure S1). Thus, LEBV, like SANTV (Félix et al. 2011), could not infect *C. elegans*, at least in these multigenerational assays relying on the whole viral cycle.

### Laboratory infection of *C. briggsae* natural isolates reveals variation in sensitivity to SANTV and LEBV

We further focused on the intraspecific variation of *C. briggsae* in infections by SANTV and LEBV. We assayed a set of 40 wild isolates of *C. briggsae* representative of genetic and geographic diversity of the species (Cutter et al. 2010; Thomas et al. 2015) for their ability to sustain infection by either virus in mono-infection experiments. These strains were inoculated with SANTV or LEBV in parallel (Figure 2A). After 8 days at 23°C (ca. two transfers and three host generations, requiring the whole viral cycle of horizontal transmission among animals of different generations), we scored the proportion of infected *C. briggsae* animals by FISH, taken here as a proxy for viral sensitivity (Table S4). Figure 2B shows the grand mean proportion of infected animals across three independent assays, with two or three replicate infections per assay and 100 scored individuals per replicate (design in Figures 2A and S2A). Many *C. briggsae* strains were sensitive to both viruses or resistant to both. Overall, we found a significant correlation among *C. briggsae* isolates in their sensitivity to SANTV and LEBV (Figure 2C). However, specificity could be found, particularly for the strains HK104 and HK105 that were fully resistant to LEBV yet highly sensitive to SANTV. Conversely, a few strains such as JU2160 were specifically resistant to SANTV but sensitive to LEBV. We confirmed the FISH results by assaying the viral load by RT-qPCR in similar infection experiments on a subset of strains (Figure S2A). The fact that we found specificity in both directions implies an interaction between host isolate and virus species.

Nevertheless, to confirm that the strong pattern of specificity in strains such as HK104 was not the consequence of a different potency of the initial LEBV and SANTV inoculates, we used serial dilutions of preparations of each virus on selected strains and assayed infection, either by FISH or by RT-qPCR in two separate experiments (Figure S3). Overcoming the difficulty in comparing viral preparations (see Frézal et al. 2019), the serial dilution results showed that the *C. briggsae* strains differed in their ability to be initially infected by a given amount of viral inoculate, as well as by the proportion of infected animals after 7-8 days. For example, *C. briggsae* strain JU516 required a higher initial viral concentration than JU1264, especially for SANTV. Importantly, these experiments clearly showed the specificity of infection of the HK104 strain by LEBV, compared to JU516 or JU2160. Using the SANTV variant JUv1993 (Frézal et al. 2019), we could also observe some infection of strain JU1377 or rarely AF16, suggesting that sensitivity of the host may depend on the viral genotype. In addition, when assaying at an earlier timepoint (3 days post-infection), infection of JU1399 and ED3032 mostly occurred in adults of the first generation (60% and 42% FISH-positive animals, respectively), raising the possibility that these strains were competent for viral entry and replication but defective in the production of new infective virions.

The proportion of infected animals is a quantitative trait, but for the sake of simplicity, we colored the strains in Figures 2C and 3 using four categories based on a binary score for each virus: resistant to both viruses (orange); sensitive to both (green); specifically resistant to LEBV (light blue); specifically resistant to SANTV (purple) (keeping some strains with an unclear status in black). In *C. briggsae,* the strong population genomic structure was shown to match geography (Cutter et al. 2006; Cutter et al. 2010; Thomas et al. 2015). Most of the strains from the temperate climates were found to be sensitive to both viruses; this set included JU1264 and JU1498, the two *C. briggsae* strains in which the viruses were first discovered. In contrast, most strains of tropical origin, such as the reference strain AF16, were resistant to both viruses. The most interesting strains were HK104 and HK105 which were specifically resistant to LEBV but highly sensitive to SANTV. Conversely, a few strains such as JU2160 from Zanzibar were specifically resistant to SANTV but sensitive to LEBV.

We thereafter focused on AF16, HK104 and JU1264 (or JU1498, close genetic relatives from France; Figure 3) as representatives of resistance, specificity and sensitivity, respectively.

### Sensitivity to the *C. briggsae* viruses does not correspond to a defective small RNA response in the host

To test whether the viruses elicited a small RNA interference response in *C. briggsae* as was observed in *C. elegans* with the Orsay virus, we infected *C. briggsae* AF16, HK104 and JU1264 isolates with SANTV or LEBV during a single generation and sequenced their small RNA content (design in Figure S5). We performed mono-infections by a single virus or co-infection of SANTV JUv1264 and LEBV JUv1498 (or of SANTV JUv1264 and SANTV JUv1993). We used a phosphatase treatment that enabled the detection of 1° siRNAs as well as 2° siRNAs and mapped the sequence reads to the genome of either virus. The expectation from the Orsay virus-sensitive *C. elegans* wild isolate JU1580 was that the sensitive strains may be defective in the small RNA response: for example JU1580 carries a natural *drh-1* deletion and similar to *drh-1* mutants, mounts a weaker antiviral 2° siRNA response (small RNAs of 22 nucleotides starting with a G) (Ashe et al. 2013; Coffman et al. 2017). Recent studies using *C. briggsae* AF16 and HK104 demonstrated an endogenous small RNA response with 1° and 2° siRNAs with the same characteristics as in *C. elegans* N2 (Fusca et al. 2022), but specific responses to these exogenous viruses have not been studied.

In the sensitive *C. briggsae* JU1264, both viruses replicated at high levels and viRNAs were detected against both viruses (Figure 4A, Table S5), including both sense 1° siRNAs and antisense 2° 22G-siRNAs (Figure 2B, Table S5). This demonstrated the ability of JU1264 to mount a proper RNA-directed antiviral immune response yet this small RNA response was insufficient to prevent viral propagation.

**Figure 4.**
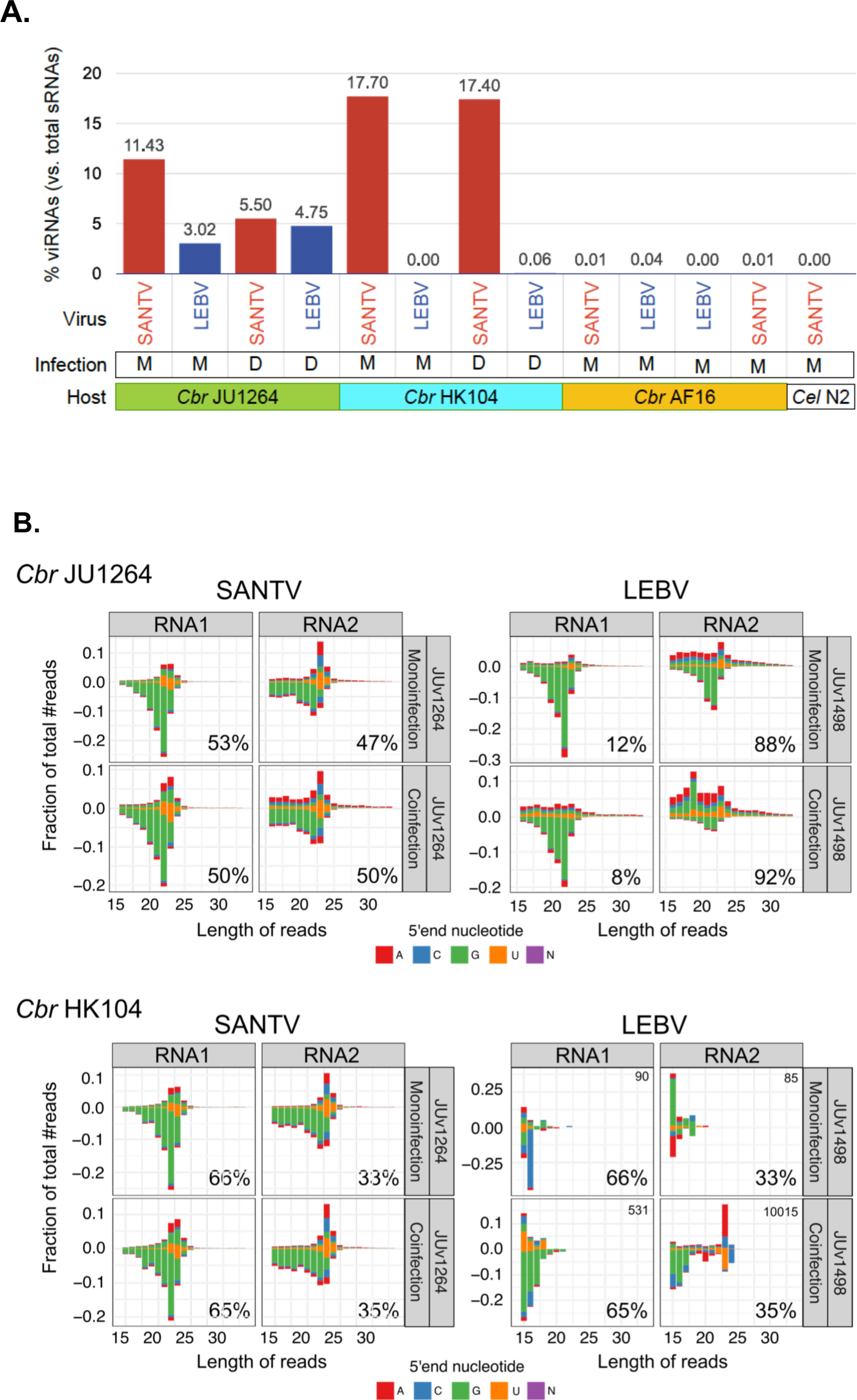
Differential pattern of small RNAs against SANTV and LEBV. (A) Proportions of 16-33 nucleotide reads mapping to viral genomes in infection experiments performed in parallel in three *C. briggsae* strains and a *C. elegans* strain. The letters M and D refer to mono- and co-infections with SANTV-JUv1264 and LEBV-JUv1498 viruses. The *C. elegans* N2 infection is used as a control for reads from the viral inoculum. Uninfected *C. briggsae* AF16 animals were used as a negative control. The host strain names are color coded as in Figure 3, and viruses are color coded as in Figure 1. (B) Differential pattern of small RNAs mapping to the two RNA molecules of the two viruses in two *C. briggsae* hosts, JU1264 and HK104. The stack bar charts show the distribution in length and 5′ nucleotide of small RNAs mapping onto each viral RNA. Negative values correspond to antisense small RNAs. The percentage of small RNAs mapping to the RNA1 and RNA2 molecules normalized to SANTV RNA2 length were computed for each infection condition and indicated on the bottom right of each graph. See Tables S4 and S5 for detailed counts and Figure S6 for mapping along the viral genomes.

In the resistant *C. briggsae* AF16, viral small RNAs (viRNAs) were absent (Figure 4A, Table S5), suggesting that neither LEBV nor SANTV could enter its intestinal cells. In *C. briggsae* HK104, only SANTV replicated at high levels, and the small RNA response to this virus was similar to that in JU1264. After an inoculation of HK104 by LEBV, viral siRNAs did not accumulate, suggesting that this virus may not be able to enter or to replicate at sufficient levels (Figure 4A,B). Therefore, whereas the *C. elegans* N2 reference strain defends itself against ORV via its small RNA response (Félix et al. 2011) (Ashe et al. 2013; Shirayama et al. 2014; Coffman et al. 2017), this does not seem to be the case for the tested resistant *C. briggsae* strains, where the viruses appeared to be blocked at entry or at early steps of the viral cycle.

### The two viral RNAs of SANTV and LEBV elicit different patterns of small RNA response

We further compared the small RNA responses to the two different viruses. In *C. briggsae* JU1264 where they both thrived, SANTV elicited a stronger anti-sense response (minus strand) with an enrichment at 22 nucleotides starting with a G (hallmark of 2° siRNAs). In comparison, LEBV infection resulted in more abundant sense RNAs of various sizes (viral genome degradation products) and a peak at 23 nucleotides on the sense strand (hallmark of 1° siRNAs) (Figure 4B).

The RNA1 molecule of both viruses triggered a marked 2° siRNA response in the JU1264 strain (and in HK104 for SANTV) (Figure 4B, Table S5). What differed between the two viruses was the proportion of reads mapping to RNA1 versus RNA2. While this proportion was similar for SANTV in JU1264, more reads mapped to LEBV RNA2 than to RNA1 (Figure 4B). LEBV RNA2 was highly degraded in small RNAs of various sizes on the sense strand, while SANTV RNA2 appeared degraded with the characteristic length of 1° siRNAs at 23 nucleotides. This differential pattern between RNA molecules and viruses was even more striking upon co-infection by both viruses – as if SANTV RNA1 was particularly targeted by the siRNA machinery. The distribution was similar for the SANTV variants JUv1264 and JUv1993 (Table S5). The proportion of reads mapping to RNA1 was even higher in the *C. briggsae* HK104 strain than in JU1264 (Figure 4B).

The distribution of small RNAs along the viral genome differed between SANTV and LEBV: SANTV displayed an enrichment in the first 200 bp at the 5’ end of RNA1 in both *C. briggsae* JU1264 and HK104 strains, which was not the case for RNA2 or either of LEBV RNAs in JU1264 (Figure S6).

### Crosses between AF16 and HK104 indicate two major QTLs on chromosomes IV and III

For genetic studies on the host side, we focused on the *C. briggsae* HK104 strain, which was specifically sensitive to SANTV and resistant to LEBV. The HK104 strain had been used previously by others as a source of polymorphic genetic markers compared to the reference strain AF16 (Fodor et al. 1983; Baird et al. 2005; Hillier et al. 2007; Inoue et al. 2007; Koboldt et al. 2010). Advanced Intercross Recombinant Inbred Lines (AIRILs) between AF16 and HK104 had been built and genotyped, in order to advance *C. briggsae* genetics and genomics (Ross et al. 2011). These recombinant lines allowed us to assess the genetic architecture of the difference in SANTV sensitivity between AF16 (resistant) and HK104 (specifically sensitive to SANTV). Figure 5A shows the distribution of SANTV sensitivity in 65 of these RILs between AF16 and HK104. Whereas 25 out of 65 RILs (38%) showed full resistance, similar to the AF16 parent, the remaining lines displayed a wide range of percentages of infected animals.

**Figure 5.**
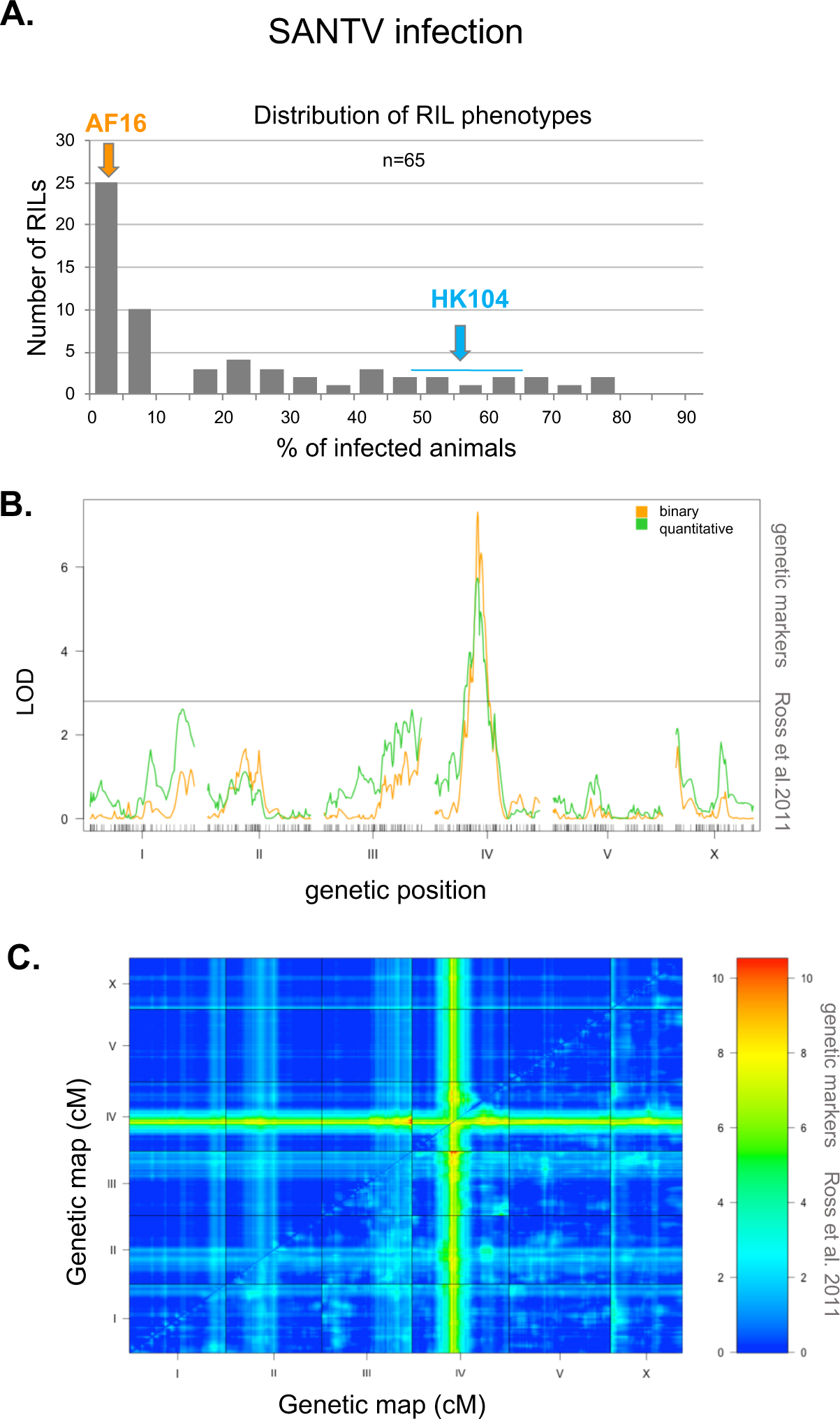
A major locus on chromosome IV underlies the variation in SANTV infection rate between *C. briggsae* AF16 and HK104. (A) Distribution of the proportion of infected animals after exposure to SANTV in Advanced Recombinant Inbred Lines (RILs) between AF16 and HK104. The mean phenotypes of the parents are shown above the graph. See Table S6 for detailed data. (B) Quantitative genetic mapping of the proportion of infected animals after infection with SANTV. Green line: phenotype coded as a quantitative trait. Orange line: phenotype coded as binary. The genetic map with the markers along the six chromosomes is shown on the *x* axis. The black horizontal line denotes the *p*<0.05 significance threshold calculated using 10,000 permutations of the data. The QTL peak on chromosome IV is at 77.2 cM. (C) LOD score grid for the two-QTL analysis, represented with a color scale from blue=0 to red=10. The upper left triangle corresponds to the additive model and the lower right triangle to the full model. Both analyses point to the same significant regions, i.e. a main locus on chromosome IV and a second one on the right tip of chromosome III.

**Figure 6.**
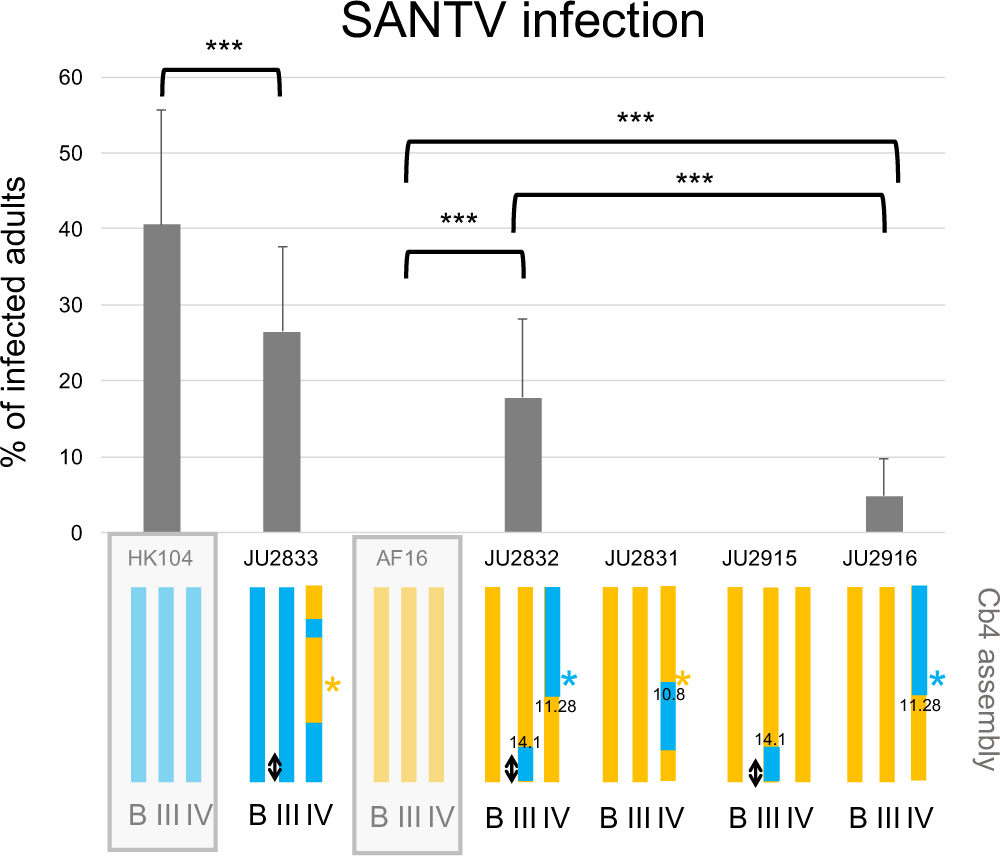
Confirmation of the AF16 x HK104 QTLs on chromosomes IV and III using Near Isogenic Lines (NILs) The genotypes of the lines are shown schematically below the graph, with orange representing the AF16 background (resistant) and green the HK104 background (sensitive to SANTV). The parental backgrounds are highlighted by a light grey rectangle. The color of the star sign at the QTL position on chromosome IV represents the inferred allelic state in the corresponding line. The QTL on chromosome III is represented by a double arrow. “B”: background (the other chromosomes). Positions of recombination breakpoints are in megabases (Mb). Detailed genotypes can be found in Table S10. The infections by SANTV were performed on two different days, with two replicates each day and 100 individuals per replicate. Bars represent the standard deviation among replicates. ***: *p*<0.001 in a generalized linear model (glm) taking day and replicate into account, each NIL being compared to its relevant background parent.

We performed a QTL analysis using the known genotypes of these RILs from Ross et al. (2011). The resulting scores from the one-QTL analysis are shown in Figure 5B, plotted using the infection data as a binary trait (presence or absence of infected animals) or as a quantitative trait (proportion of infected animals, the mean of two infection experiments) (Table S6). A single region on chromosome IV displayed LOD scores above the threshold, meaning that genetic variation in this region explained a part of the phenotypic variation. The peak on chromosome IV is located around 77.3 cM of the genetic map (Ross et al. 2011). In addition to this main QTL, two regions were close to the threshold in the quantitative trait analysis (green curve in Figure 5B). We further performed a two-QTL analysis testing every pair of positions along the genome. Figure 5C shows the LOD (Logarithm of the ODds) score for the additive and interaction models of the two-QTL analysis. Both indicated a main locus on chromosome IV and a second locus on the right tip of chromosome III.

### Near Isogenic Lines (NILs) confirm both QTLs between AF16 and HK104

To test the QTLs and quantify their effect on the phenotype, we created near isogenic lines (NILs) by backcrossing the QTL regions of chromosomes III and IV of one parent into the other parental background. We obtained the following NILs. JU2831 carried the AF16 background with an introgression of the 11-14.3 Mb chromosome IV region from HK104 (Figure 6 and Table S6). JU2832 carried two introgressed segments from HK104 in the AF16 background: 0-11.28 Mb on chromosome IV and 14.1 Mb-right tip on chromosome III. JU2915 and JU2916 corresponded to the single introgression from JU2832 from chromosomes III and IV, respectively. Conversely, JU2833 carried the chromosome IV introgressions (0-2 and 6-13.8 Mb) from AF16 into the HK104 background (Figure 6).

The SANTV sensitivity levels of these introgression lines are shown in Figure 6. JU2832, bearing the HK104 chromosomes III and IV QTL alleles in the resistant AF16 background, was sensitive to SANTV. JU2916 with the HK104 QTL region on chromosome IV was also sensitive to the virus, but in a significantly lower proportion. In contrast, JU2915 which only had the HK104 QTL region on chromosome III was resistant to SANTV (note however that we observed some infected animals in other replicates, whereas no infected AF16 animal was observed in parallel experiments). These results confirmed the QTLs on chromosomes IV and III, with the weaker chromosome III QTL requiring the presence of the HK104 allele on chromosome IV for expression in the AF16 background. Conversely, in the HK104 background, the QTL region of AF16 on chromosome IV in strain JU2833 lowered the proportion of infected animals compared to HK104, but did not abolish it, confirming the importance of other genomic regions. Finally, the JU2831 strain was fully resistant, which allowed to narrow down the QTL region between the positions at ca. 10,000,000 and 10,789,370 bp on chromosome IV (coordinates on C*. briggsae* AF16 Cb4 assembly).

We further tested whether the QTL affected LEBV sensitivity and found that the NILs JU2832 and JU2833 were fully resistant to LEBV (n= 100 animals; a positive control JU1498 was infected). The QTLs thus appear to specifically affect the interaction with SANTV in this context.

The major QTL region on chromosome IV contains many polymorphisms. Among them, two potential candidates are 14 non-synonymous polymorphisms in the *Cbr-rsd-2 (CBG01755)* gene and one in a paralog of *C. elegans rde-11 (CBG01824)*, since *C. elegans rsd-2* and *rde-11* mutations render the animals sensitive to the Orsay virus (Guo et al. 2013 and Table S7). A replacement by CRISPR/Cas9 genome editing of the non-synonymous polymorphism in *CBG01824* coupled with a deletion in the *Cbr-rsd-2* ortholog did not render *C. briggsae* AF16 sensitive to SANTV (Table S7). Further work is needed to identify the causal polymorphism(s).

### Recombinant Inbred Lines (RILs) between JU1498 and HK104 indicate a main QTL on chromosome II

We further built RILs after a cross between HK104 (resistant to LEBV) and JU1498 (sensitive to both viruses) and phenotyped them using LEBV infection. The phenotypic distribution of the 79 lines is shown in Figure 7A (details in Table S8). We selected at the two ends of the distribution 37 resistant lines and 23 highly sensitive lines and sequenced their genome in two pools, as well as that of each parent. In Figure 7B, we plotted the frequency of the HK104 allele along the genome for each pool (numbers in Table S9).

**Figure 7.**
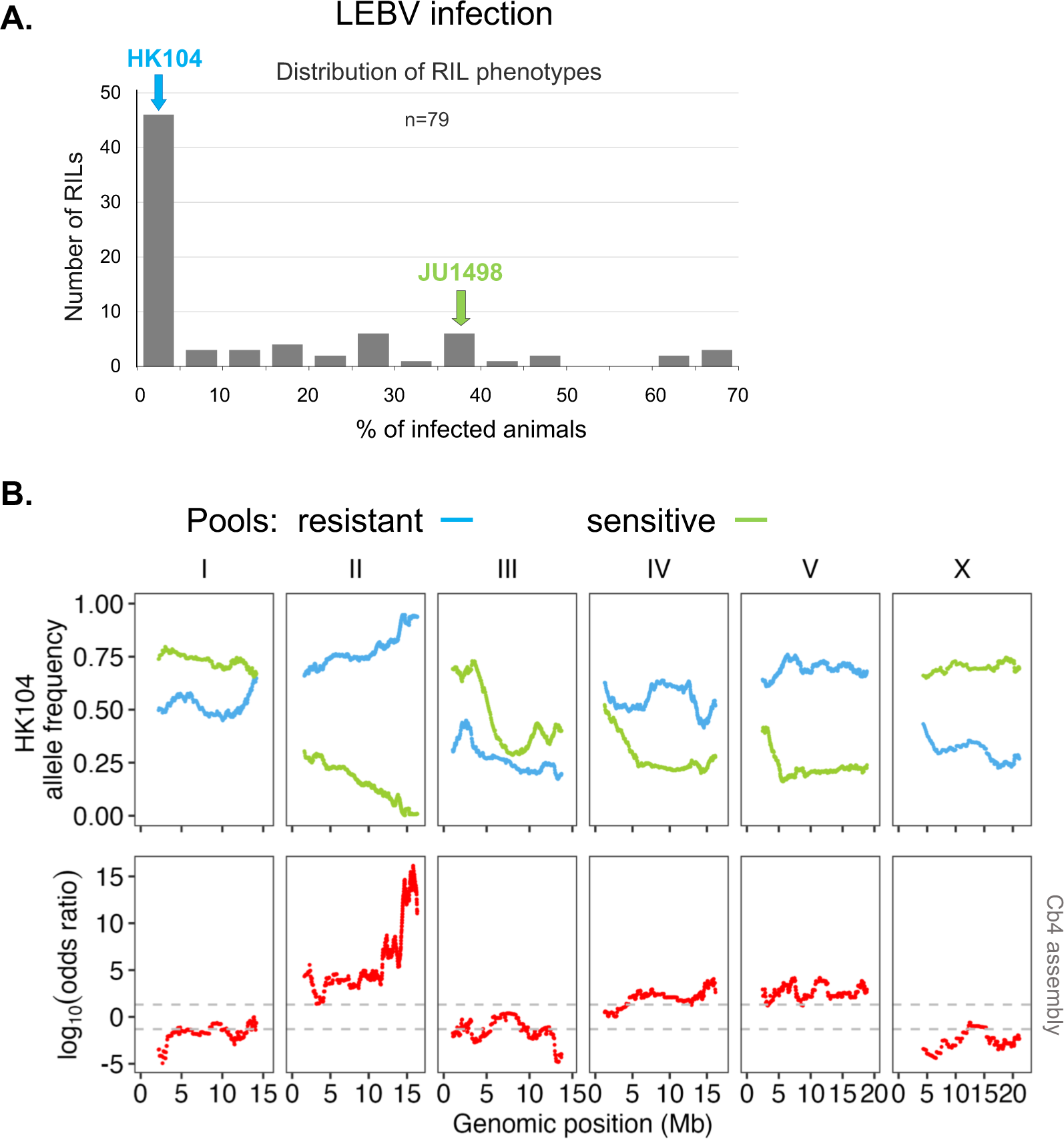
A major locus on chromosome II underlies the variation in LEBV infection rate between *C. briggsae* JU1498 and HK104. (A) Distribution of the proportion of infected animals after exposure to LEBV in Recombinant Inbred Lines (RILs) between JU1498 and HK104 (Table S8 for detailed data). (B) Bulk sequencing of pools of resistant and sensitive lines. The top plot shows the HK104 allele frequency of each pool along the *C. briggsae* genome (Cb4 assembly) (Table S9 for detailed data). The bottom plots show the LOD scores along the physical map of the six chromosomes. The dotted lines represent the threshold of significance at *p*<0.01.

We then derived a score as in Frézal et al. (2018), based on the number of lines in each pool. A highly significant QTL was found on the right tip of chromosome II, as well as possibly minor ones, including a possible transgressive QTL on chromosome X, for which the HK104 allele appeared to confer higher sensitivity.

### Near Isogenic Lines (NILs) confirm the QTL between JU1498 and HK104

To directly test the effect of the main QTL on chromosome II, we introgressed this region of one parent into the other parental background. All introgression lines JU3244-3247 obtained by backcrossing chromosome II of the LEBV-sensitive strain JU1498 into the background of the resistant strain HK104 enabled LEBV infection (Figure 8). Conversely, the reciprocal introgression line JU3241 was fully resistant to LEBV. Similar to both parental lines JU1498 and HK104, these strains were sensitive to SANTV. These experiments confirm that a major locus on the right part of chromosome II is responsible for a large part of the difference in LEBV sensitivity between JU1498 and HK104.

**Figure 8.**
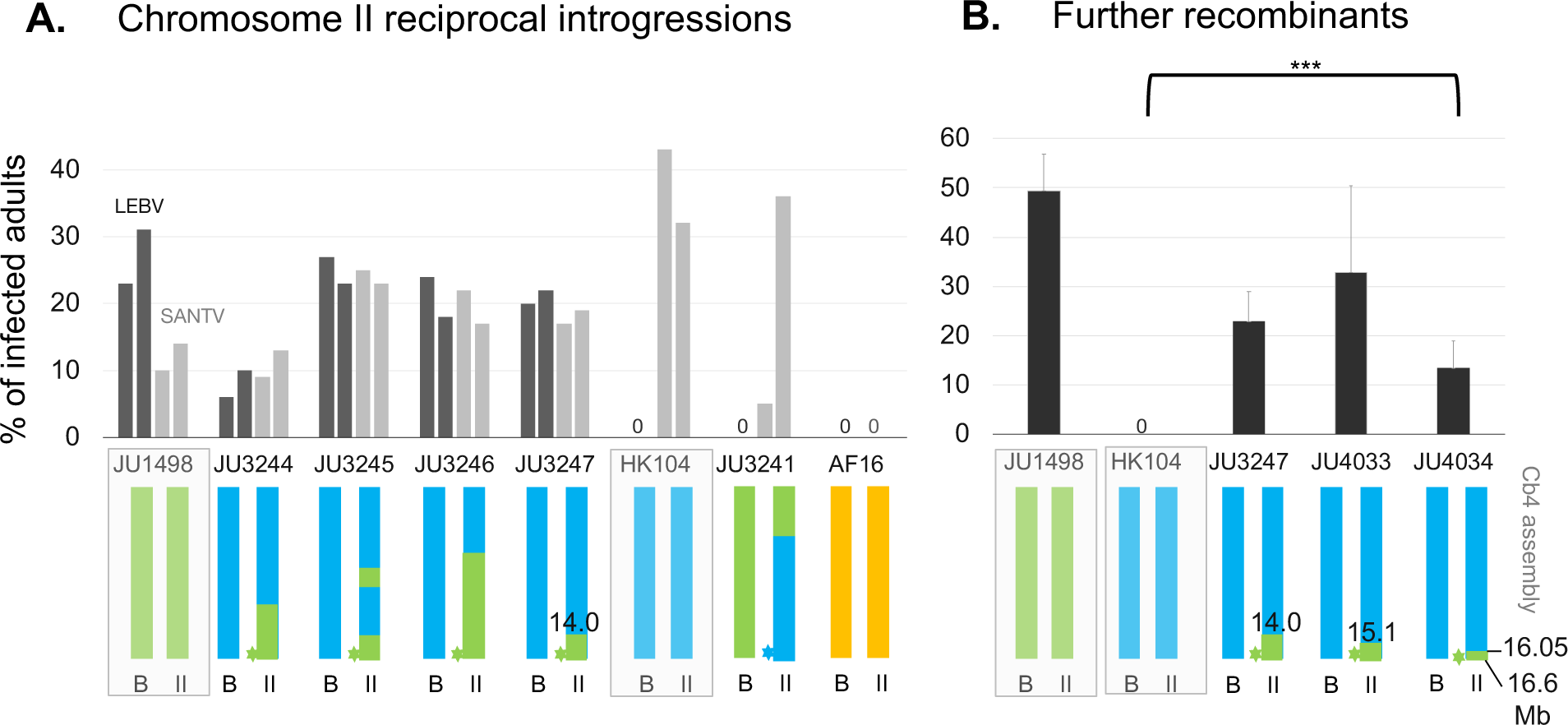
Confirmation and fine mapping of the JU1498 x HK104 locus using Near Isogenic Lines (NILs) (A) Reciprocal introgressions of the right end of chromosome II. Infections by LEBV (black) and SANTV (grey) were performed in duplicates. n=100 animals, except for SANTV infection of JU3241 (details in Table S10). The color of the star (*) at the QTL position represents the inferred state in the corresponding line. (B). Assays of further recombinants of the chromosome II QTL, showing the mean of three infection replicates. Detailed genotypes and scorings can be found in Table S10. Positions of breakpoints in Mb are indicated in relevant cases. ***: *p*<0.001; **: *p*<0.01 comparing strains pairwise using a generalized linear model.

We screened for recombinants in the QTL region after a further cross of strain JU3247 with HK104. We found that the rightmost tip of chromosome II starting at 16.05 Mb in strain JU4034 was sufficient to confer LEBV sensitivity in the HK104 background. This restricted the QTL interval to II:16.05 - 16.6 Mb (coordinates of Cb4 assembly).

## Discussion

We previously reported on the diversity of *Caenorhabditis* noda-like viruses (Frézal et al. 2019). Here we provided groundwork for studying on the host side variation in the interaction between *C. briggsae* and its intestinal viruses. Our results show that, although not yet receiving widespread attention, *C. briggsae* presents an excellent model for delving into host-virus specificity.

### Genetic basis of viral resistance in *C. briggsae*

We observed an overall correlation between susceptibility of different *C. briggsae* isolates to the two viruses SANTV and LEBV (Figure 2). This correlation suggests that a part of the natural genetic variation in viral susceptibility operates in a mechanism common to both viruses.

However, a few *C. briggsae* strains break this correlation in sensitivity between SANTV and LEBV, most strikingly HK104 and HK105 from Japan that are specifically resistant to LEBV (Figure 3). This specific pattern implies another resistance mechanism, specific to LEBV (or conversely, a SANTV-specific susceptibility mechanism).

We focused on strain HK104 for genetic studies using crosses to either the doubly resistant AF16 or to the doubly sensitive JU1498. The genetic loci detected in the two crosses differ (summary in Figure 9). The NILs exchanging the alleles at each QTL did not affect sensitivity to the other virus. A possibility is that the tip of chromosome II QTL encodes a specific factor necessary for LEBV entry or replication, such as a viral receptor or a host factor specifically required for LEBV translation or RdRP activity, and that *C. briggsae* HK104 is defective in this factor. Alternatively, *C. briggsae* JU1498, like many temperate strains, may have lost a LEBV-restricting factor. Finally, it is possible that this chromosome II QTL factor affects both viruses but that the specific SANTV sensitivity comes from another locus. The main QTL on chromosome IV (or that on chromosome III) in the AF16 x HK104 cross may correspond to a SANTV-specific factor. Alternatively and symmetrically, it may correspond to a general factor in a context where LEBV would be restricted by other loci in the AF16 and HK104 genomes. Molecular identification of the QTL or crosses between AF16 and JU1498 will be necessary to evaluate which scenario is most likely.

**Figure 9.**
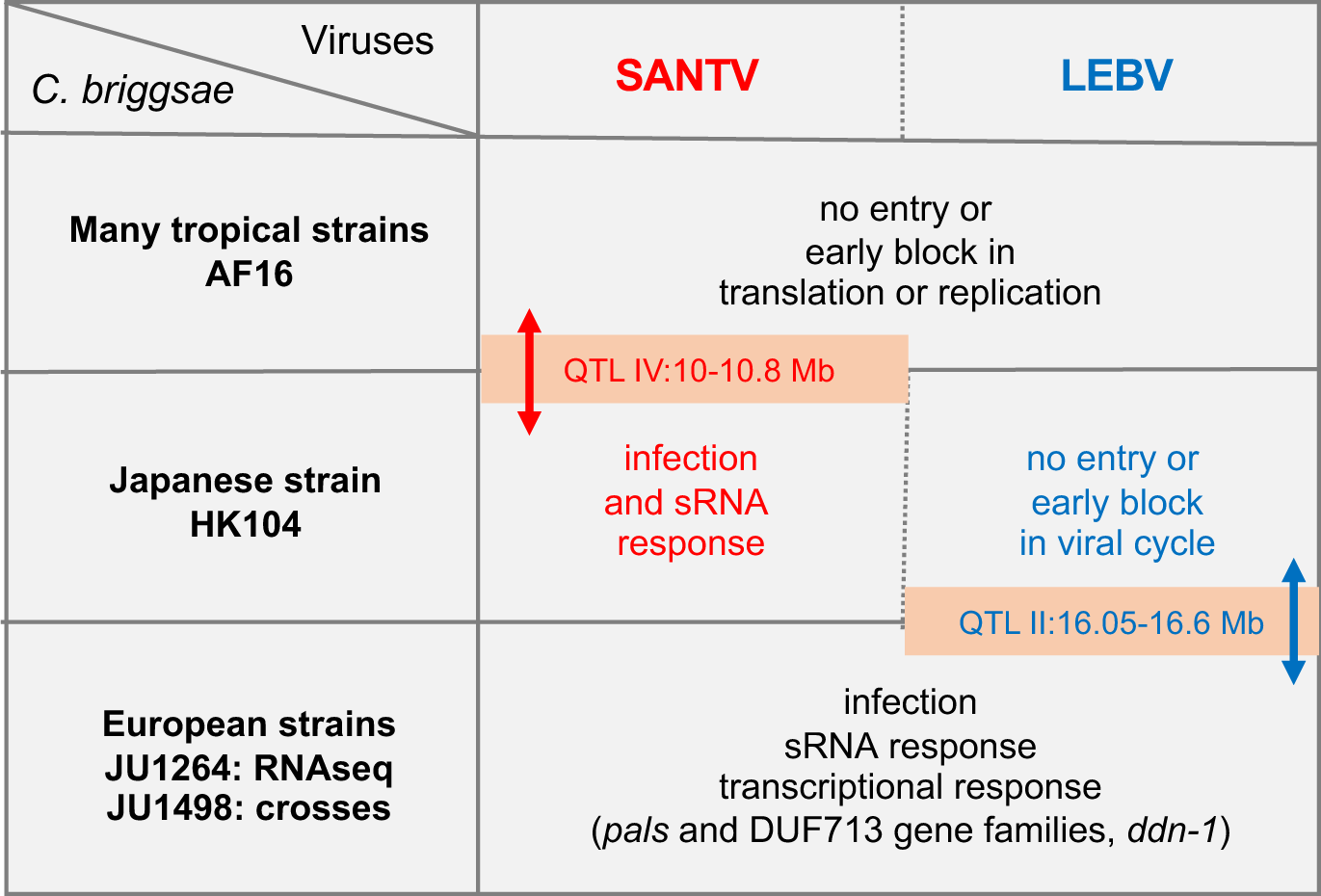
Schematic summary of the findings. The transcriptional result in *C. briggsae* JU1264 are from Chen et al. (2017).

Because the first experiment appeared positive, we assessed the *CBG01824* non-synonymous SNP replacement and deletions in seven independent infection experiments. Altogether, we noted an initial increase in viral sensitivity after CRISPR genome, which was abolished along further experiments (Table S7). This possible transient effect may parallel previous observations on CRISPR replacement assays for the mortal germline phenotype (Frézal et al. 2023). We do not know whether this may be a gene-specific or aspecific effect of oligonucleotide and/or Cas9 injections. We also tested *Cbr-rsd-2* deletions and they did not render C. *briggsae* AF16 sensitive to the viruses, further corroborating that the block in infection occurs earlier (Table S7).

The intervals still contain many polymorphisms and those tested so far were thus inconclusive. In instances where multiple loci contribute to the candidate QTL or other minor QTLs influence the phenotype, the task of identifying a candidate locus for that phenotype can become difficult (Noble et al. 2017; Bernstein et al. 2019; Sterken et al. 2021). We plan to further narrow down the intervals using CRISPR-mediated recombination (Zdraljevic et al. 2023). Better assemblies of the regions in the relevant strains will also be required.

### Small RNA and transcriptional response of *C. briggsae* to viral infections

The viRNA responses we detected in *C. briggsae* differ from those in *C. elegans* strains, whether the ORV-susceptible JU1580 (*drh-1/RIG-I* defective) or the relatively resistant N2 (*drh-1+*). On one hand, in contrast to the *C. elegans drh-1* defective wild strains, viral infection of *C. briggsae* JU1264 and HK104 occurs despite an apparently normal siRNA response (note that a high viral load corresponds to more viral RNA substrate available for cleavage). On the other hand, the reference *C. elegans* N2 strain is relatively resistant to viral infection due to a detectable small RNA response (Félix et al. 2011; Ashe et al. 2013; Sarkies et al. 2013). Instead, in the resistant *C. briggsae* strains (AF16 to both viruses; HK104 to LEBV), viRNAs are absent, suggesting an absence of receptor-mediated entry into the nematode intestinal cells or a block at an early step of the viral cycle such as translation or replication (Figure 9).

SANTV and LEBV elicited different small RNA patterns when assayed in the same susceptible host (JU1264), particularly in co-infections. However, this difference was mostly explained by the difference between RNA1 and RNA2, and the preferential amplification of 22G 2° siRNAs matching the 5’ end of SANTV RNA1. Such an enrichment at viral RNA ends was observed in *C. elegans drh-1* mutants for ORV RNA1 (5’ end) and RNA2 (both ends) (Coffman et al. 2017). A possibility, which appears to us unlikely, is that all other RNA ends are missing in the present SANTV and LEBV genome assemblies (see Ashe et al. 2013; Coffman et al. 2017). Alternatively, this SANTV RNA1 enrichment may result from 5’-end recognition, or a specific secondary/double-stranded structure that may overwhelm the small RNA machinery. Upon viral entry, RNA1 is likely the first to be translated and replicated as it encodes the viral polymerase. Whether a specific sensitivity mechanism to SANTV in *C. briggsae* HK104 is related to this differential pattern of small RNAs on the two viruses is unlikely given the absence of small RNA response to LEBV, suggesting an earlier block.

Besides this small RNA response, Chen et al. (2017) studied the transcriptional response of *C. briggsae* JU1264 to SANTV infection, and compared it with the *C. elegans* response to ORV, in the N2 reference and a *rde-1* mutant (defective in the RNA interference response). They found that orthologous genes were upregulated in *C. elegans* and *C. briggsae*, for instance: 1) specific *pals* family members (Reddy et al. 2017; Lazetic et al. 2022); 2) genes encoding DUF713 domain-containing proteins (e.g. *CBG18525* in *C. briggsae*, *B0507.6* in *C. elegans*) and their adjacent genes (*CBG18525-35* and *B0507.x* genes); 3) specific C-type lectins; 4) *ddn-1*. Many of these genes are transcriptionally activated by the ZIP-1 transcription factor (Lazetic et al. 2022) and also activated by microsporidia infection, heat or protein stress in *C. elegans* in what is known as the intracellular pathogen response (IPR) (Bakowski et al. 2014; Chen et al. 2017; Reddy et al. 2017; Sowa et al. 2020; Lazetic et al. 2023). In contrast, Chen et al. (2017) did not observe in *C. briggsae* induction of ubiquitin pathway genes (MATH domain, cullins, etc.) nor of the *eol-1* ortholog (coding a RNA decapping enzyme) upon SANTV infection in *C. briggsae* JU1264. The transcriptional response to LEBV has not been studied so far.

Given the small RNA and transcriptional responses, defects in mechanisms other than small RNAs, such as ubiquitin-mediated degradation or yet unknown pathways, may explain the viral susceptibility of *C. briggsae* wild strains. The resistant *C. briggsae* strains appear resistant for viral entry or pre-replicative steps.

### Evolution of viral susceptibility in *C. briggsae*

The majority of virus-susceptible *C. briggsae* strains were collected from temperate regions. From *C. briggsae* population genomic data, either the colonization of temperate environments is a recent event or the temperate populations lost diversity due to recent selective sweeps (Cutter et al. 2006; Cutter et al. 2010; Thomas et al. 2015). Either way, it seems plausible that viral susceptibility of *C. briggsae* appeared within the temperate population, or in a tropical subpopulation followed by migration to temperate regions. Perhaps explained by a geographical sampling bias and/or the viral sensitivity pattern, SANTV and LEBV were so far solely found in Europe (Frézal et al. 2019). Thus, co-evolution of the *C. briggsae* host and its viruses may have specifically occurred in the temperate zone. The Japanese strains HK104 and HK105 are within the temperate genetic group yet distinct from the European isolates (Figure 3). The existence of specific resistance to one virus reinforces the notion of coevolution between *C. briggsae* and its natural viruses.

Whether this co-evolution is antagonistic or mutualistic remains unclear. Infection by these viruses is deleterious in standard laboratory conditions by slowing down production and in some cases lowering brood size and longevity (Ashe et al. 2013) (Figure 1). However, in *C. elegans*, evolution occurred with the *drh-1* deletion in the unexpected direction of less pathogen-resistance. Spread of this allele via selection at a linked locus is a possibility (Ashe et al. 2013). Alternatively, the *drh-1* deletion may have been selected because of pleiotropic consequences on small RNA pathways (Sarkies et al. 2013; Mao et al. 2020). Genetic attenuation of immune pathways may be beneficial for the host. Finally, viral infections themselves may protect *Caenorhabditis* against other stresses, for example by activating the transcriptional response known as the intracellular pathogen response (IPR) (Sowa et al., 2020; Castiglioni and Elena 2023). Given the case of *C. elegans drh-1* and the *C. briggsae* intraspecific pattern of sensitivity, viral sensitivity to LEBV and SANTV may be derived in *C. briggsae*.

A likely exception to this possibly derived sensitivity is the partial resistance observed in the temperate strain JU516. This *C. briggsae* strain from France is genetically very close to JU1264, JU1498 and other sensitive European strains (Figure 3) yet less sensitive to both viruses, particularly SANTV. The close genetic relatedness between these strains may be helpful in identifying the molecular genetic basis for this difference.

Exploring the factors leading to the sensitivity of potential host strains to a virus remains an open avenue for investigation, including identifying the receptors responsible for viral entry into the intestinal cells. The source of specificity could in addition be sought on the virus side, for example by reconstituting the two viruses using transgenes in *C. briggsae*, as achieved for ORV in *C. elegans* (Jiang et al. 2014). The variants in the QTL regions we identified are poised for further exploration to pin down the causal molecular variants affecting virus pathogenesis in *C. briggsae*. With the LEBV-specific *C. briggsae* JU2160 from Zanzibar or JU516 from France, we also opened the way for further studies of genetic variation in viral sensitivity in *C. briggsae*.

## Abbreviations

SANTV: Santeuil virus
LEBV: Le Blanc virus
ORV: Orsay virus
QTL: quantitative trait locus
RdRP: RNA-dependent RNA polymerase
IPR: Intracellular pathogen response
FISH: fluorescent in situ hybridization
RT-qPCR: reverse transcription followed by quantitative polymerase chain reaction
SNP: single-nucleotide polymorphism
LOD: logarithm of the odds
RIL: recombinant inbred line
NIL: near isogenic line.

## Acknowledgements

We thank Tony Bélicard for initial work on the viruses and help with the life history assays, Daria Martynow for technical help with the *C. briggsae* isolates and Amhed Vargas Velazquez with help with initial genomic analyses. We are grateful to David Wang for communication of unpublished *C. briggsae* and viral genome sequences and to Asher Cutter for sending us the AI-RILs. We thank the IBENS sequencing facility, especially Corinne Blugeon. We thank Eric Miska for discussions and Ruben Gonzalez for reading the manuscript. MAF is grateful for the hospitality of her parents in Le Blanc, including during writing of this manuscript. This work was supported by grants from the Agence Nationale pour le Recherche ANR-11-BSV3-013 and from the Fondation pour la Recherche Médicale DEQ20150331704 to MAF and ARF202209015859 to GZ. We thank Wormbase and CaeNDR. Some strains were provided by the CGC, which is funded by NIH Office of Research Infrastructure Programs (P40 OD010440).

## Supplemental information

**Figure S1.**
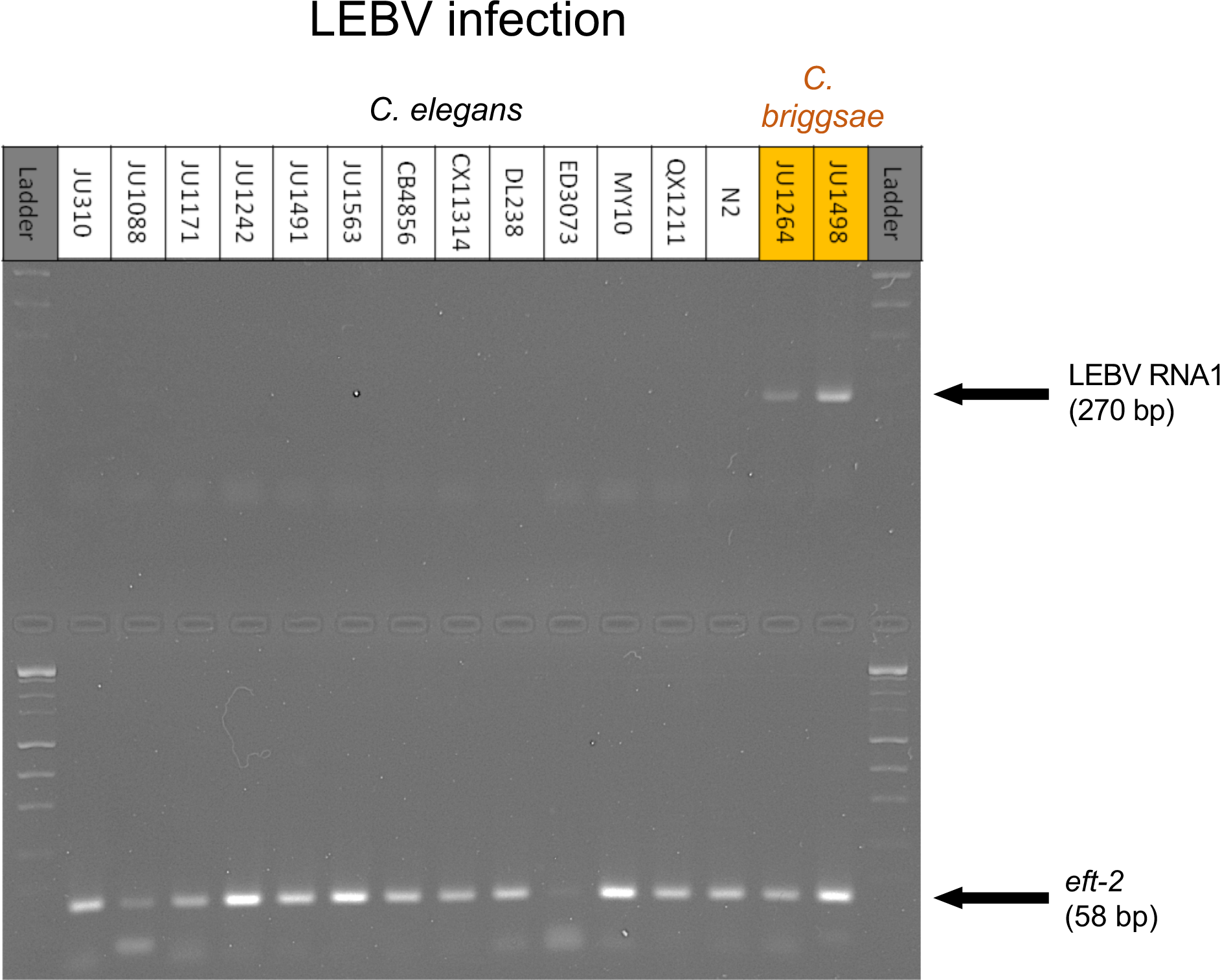
Species specificity: LEBV does not infect *C. elegans* strains. Electrophoresis on an agarose gel showing the results of a RT-PCR for LEBV RNA1. The inoculated strains are indicated on top of the gel. The infection was performed as in Figure 2A. The *C. elegans* strains include some that are infected at high levels by the Orsay virus, such as JU1491, JU1563, DL238, JU1242. A positive control for the LEBV inoculate is shown with the two *C. briggsae* strains JU1264 and JU1498. A positive RT-PCR control with *eft-2* is shown on the bottom gel.

**Figure S2.**
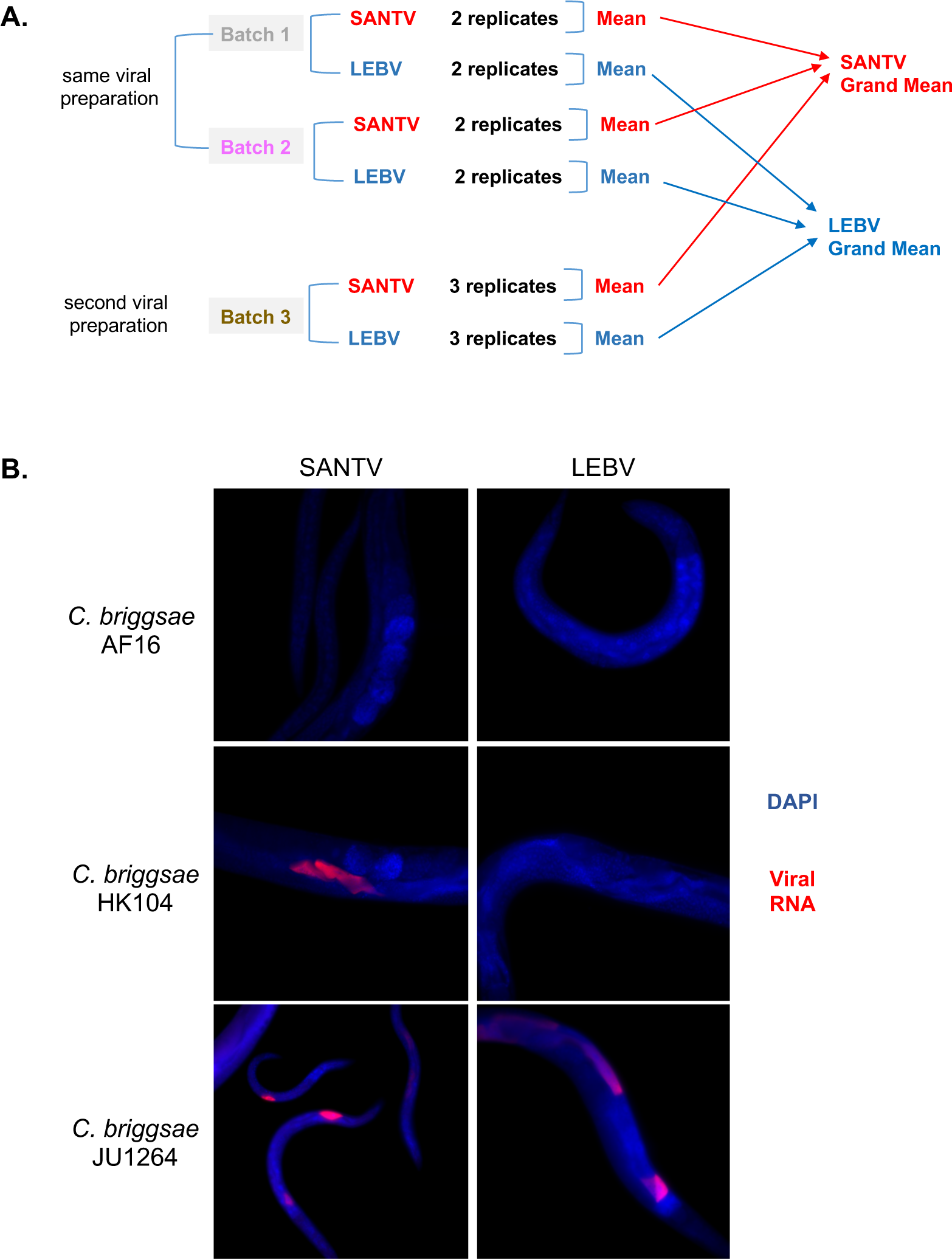
Viral infections of *C. briggsae* isolates assayed by RNA FISH for the viruses. (A) Design for the assays of the *C. briggsae* wild isolates. The data for each replicate (cf. Methods) are shown in Table S3 and the grand mean in Figure 2. The absolute level of infection differs among batches but the results are generally consistent. (B) Representative FISH images of the viruses, here using RNA1 probes for each virus. The images were acquired in the DAPI and FISH channels using a 40x objective and super-imposed in false colors.

**Figure S3.**
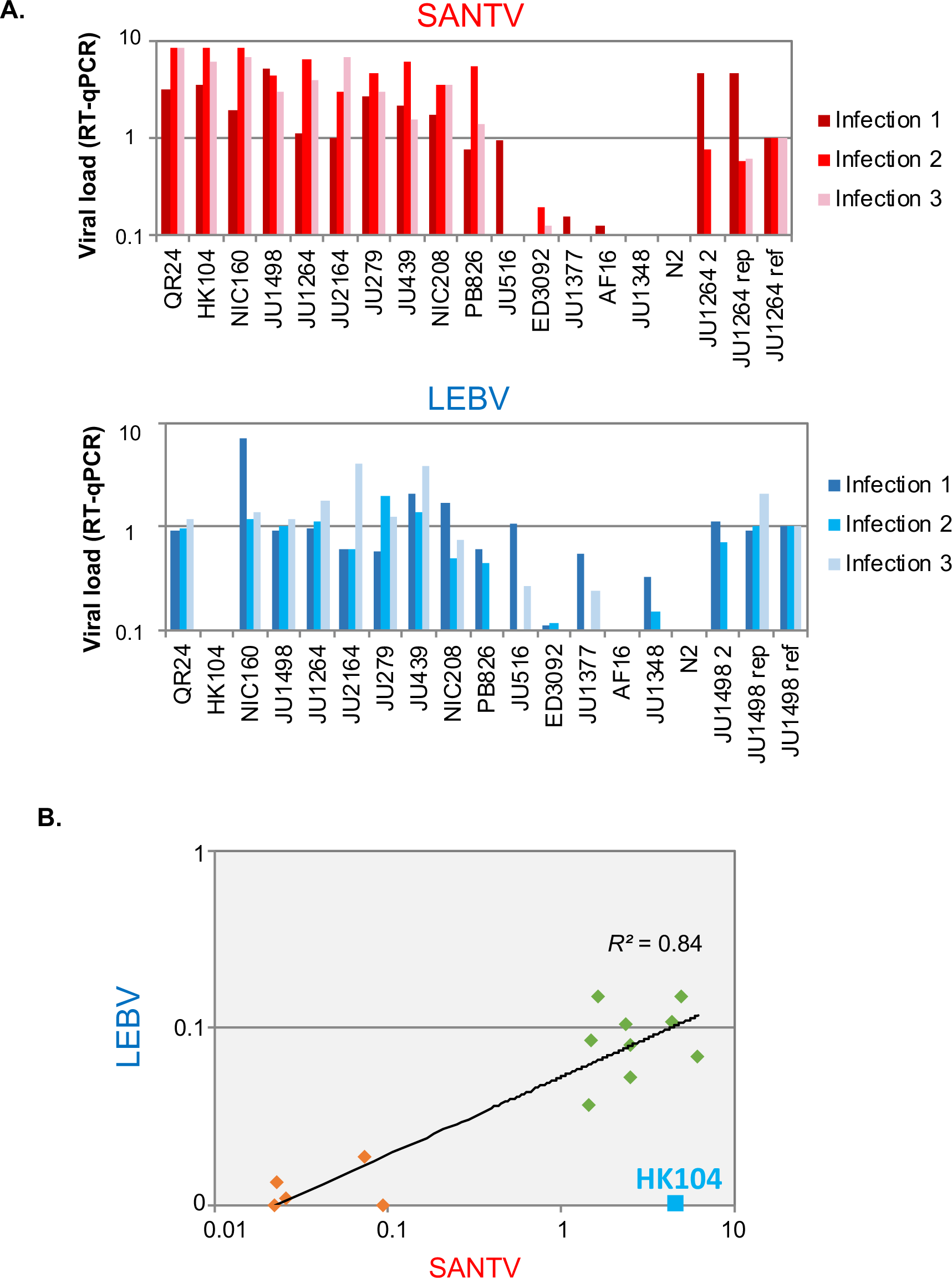
Viral infections assayed by RT-qPCR for a viral RNA. (A) Viral infections of a subset of C*. briggsae* isolates, with the viral load assayed by RT-qPCR for RNA1 of the corresponding virus. Experiments using RNA2 primers gave similar results. *C. elegans* N2 is a negative control for the initial viral inoculum. The plotted viral load corresponds to the ratio between the amplification of viral RNA1 over that of *eft-2*, normalized by that in the reference infection (JU1264 for SANTV, JU1498 for LEBV). The three infection replicates were performed in parallel. (B) Two-dimensional plot displaying the mean between the three replicates. HK104 is an outlier. The other strains show a good correlation between their sensitivity to SANTV and LEBV (regression line and correlation excluding here HK104). A larger set of *C. briggsae* isolates was assayed by FISH in Figure 2.

**Figure S4.**
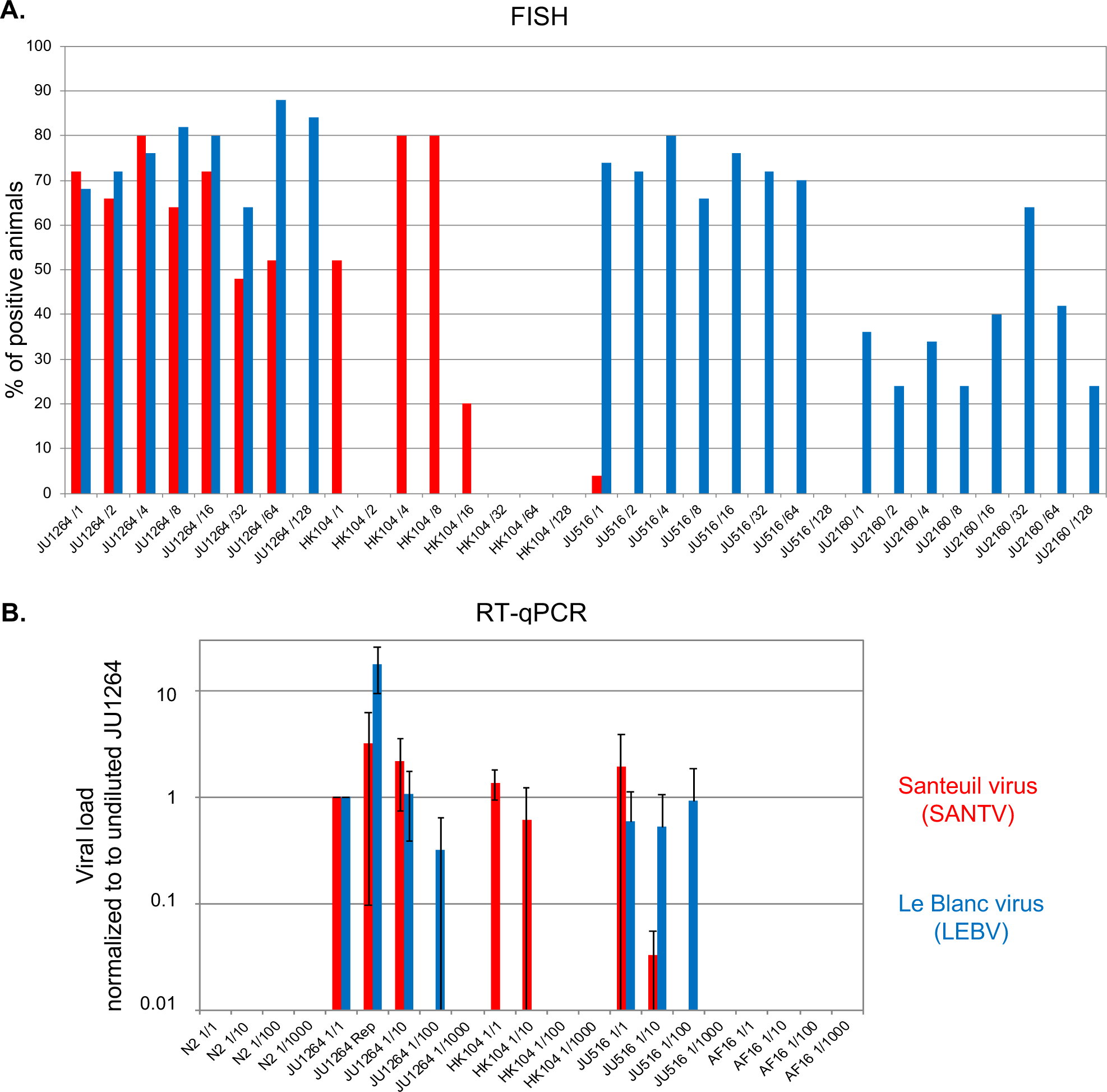
Quantitative variation in sensitivity of *C. briggsae* isolates upon serial dilutions of viral preparations. In both panels, SANTV infections are shown in red, LEBV infections in blue. (A) Proportion of infected animals as assayed by FISH staining. Four *C. briggsae* isolates were infected with successive two-fold dilutions of the same viral preparations of either SANTV JUv1264 or LEBV JUv1498, using RNA1 probes for each virus. 50 animals were scored per condition. The infection was initiated by inoculating a plate containing 5 L4 larvae and transferring 20 L4 larvae of the F1 generation, and assaying adults 3 days later, at 23°C. (B) Viral load assayed using RT-qPCR, in a separate infection experiment. *C. elegans* N2 is used as a control for amplification of the viral inoculum. The undiluted viral preparations on JU1264 are used to normalize. A separate replicate was performed and indicated as “Rep”. In each condition, n=2 RT-qPCR replicates for SANTV, 3 replicates for LEBV. Bars show standard deviation.

**Figure S5.**
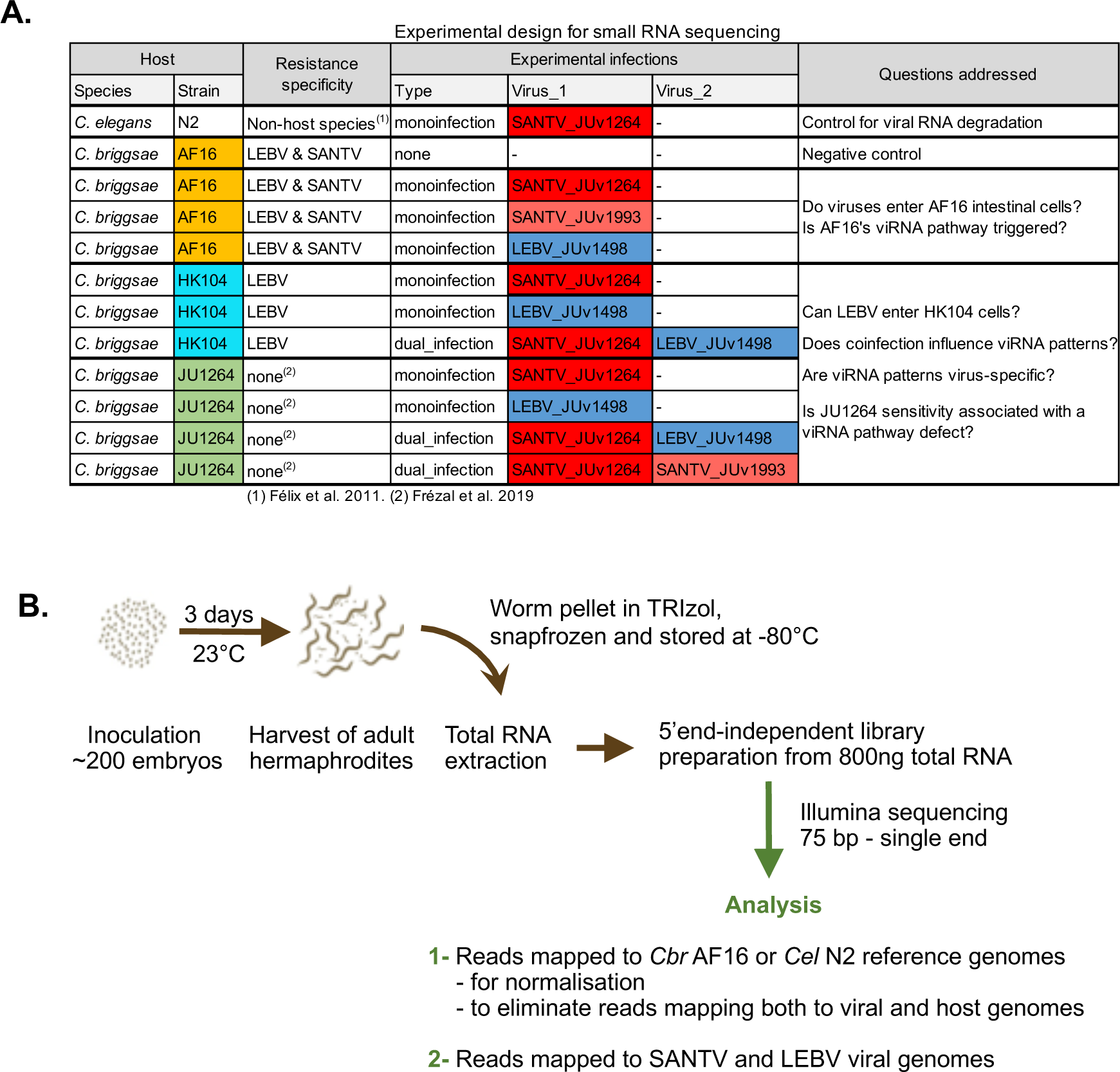
Experimental scheme to investigate the RNA-directed antiviral immune response of *C. briggsae* when infected with SANTV and/or LEBV. (A) Details of experimental conditions and questions addressed. The strain names are color coded as in Figure 3; the virus names are color coded as in Figure 1. The SANTV variant JUv1993 was also used; this variant tends to outcompete JUv1264 in long co-infection experiments (Frézal et al. 2019). (B) Schematic overview of the experimental flow.

**Figure S6.**
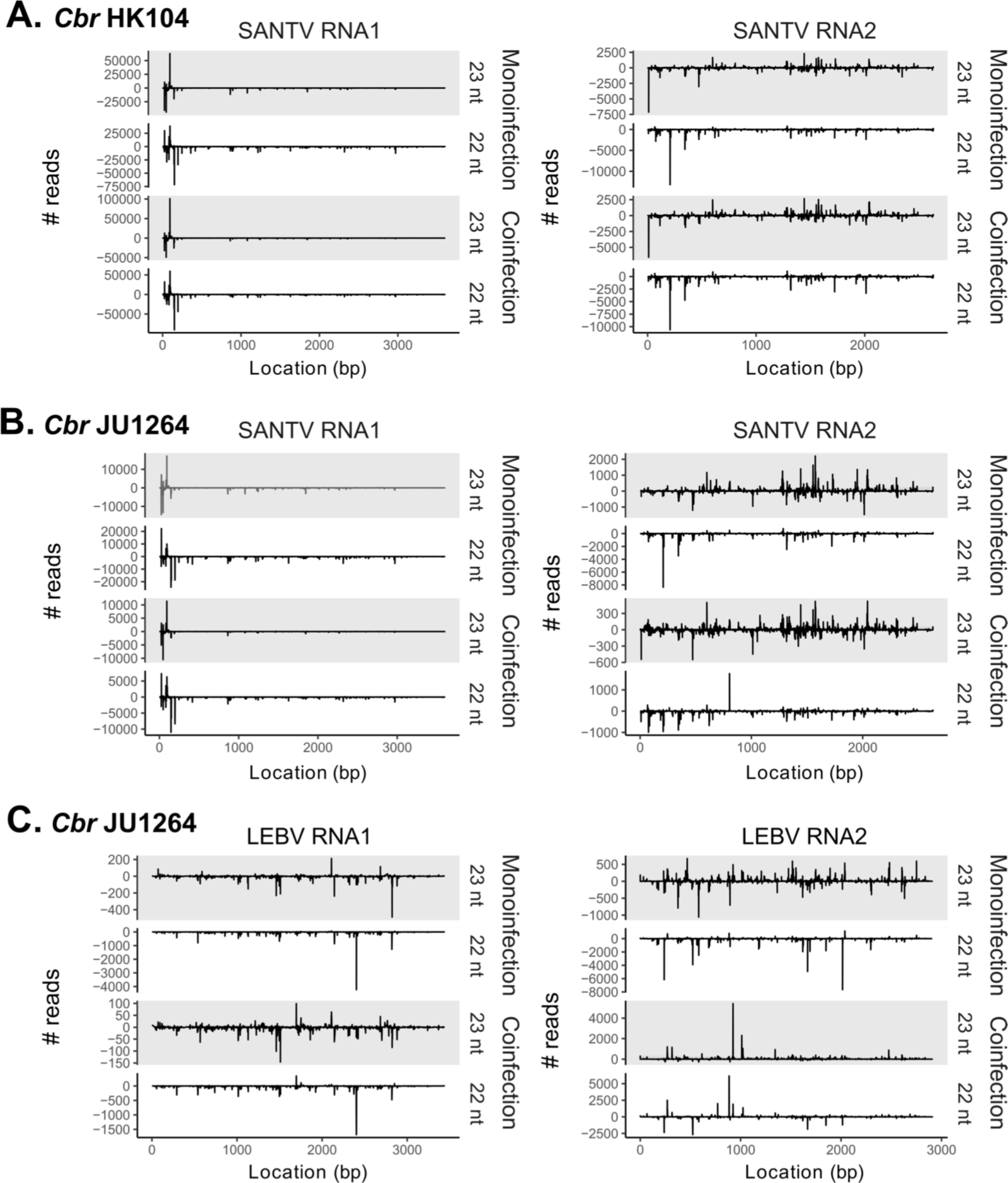
Distribution of small RNA along the viral genomes for 23 nt and 22 nt sRNAs, for each viral RNA and strand. (A) *C. briggsae* HK104 infected with SANTV JUv1264. (B) *C. briggsae* JU1264 infected with SANTV JUv1264. (C) C*. briggsae* JU1264 infected with LEBV JUv1498. Note that the different graphs of small RNA distribution along the viral genome are not at the same scale. Because of the large number of reads mapping at the 5’ end of SANTV RNA1, the other reads along the molecule are here difficult to see.

**Table S1. Strains used in this study**, including wild isolates, RILs, NILs.

**Table S2. Oligonucleotide sequences.** Different sheets indicate RT-PCR primers, pyrosequencing primers for each cross of host strains and PCR primers to genotype indel polymorphisms.

**Table S3. Raw data for survival and progeny production of infected *C. briggsae* animals.** These data correspond to the experiments in Figure 1.

**Table S4. Sensitivity of different *C. briggsae* wild isolates to the Santeuil and Le Blanc viruses.** (A,B) The results are expressed as % of infected animals as assayed by FISH for the respective virus. The two viruses were inoculated in parallel. The results are shown for the SANTV infections in (A) and LEBV infection in (B). The grand mean is shown in Figure 2. See Methods and Figure S2. (C) Serial dilutions of the viruses and scoring of infection by FISH for the virus. The results are expressed as % of infected animals. (D) Serial dilutions of the viruses and scoring of viral load by RT-qPCR.

**Table S5. Small RNA counts in viral infection experiments.** This includes control experiments, infection by SANTV variant JUv1993 (see Frézal et al. 2019) and co-infection experiments. The first sheet shows the total counts mapping to the viral and host genomes in viral infection experiments. The second sheet shows counts by categories of small RNAs of different lengths corresponding to the sense or antisense direction of the viruses. The normalized read numbers accounts for the length of the two viral RNA molecules.

**Table S6. SANTV infection of Advanced Intercross Recombinant Inbred Lines between AF16 and HK104.** The first sheet shows the infection results for two replicate infections by SANTV on the parents and the Recombinant Inbred Lines. The second sheet is the data matrix used for QTL mapping based on the mean of the two replicates, expressed as percentage of infected animals. The genotypes were taken from Ross et al. (2011).

**Table S7. CRISPR edits of the *CBG01824* and C*br-rsd-2* genes do not render *C. briggsae* AF16 sensitive to the SANTV.** The first sheet shows the candidate polymorphisms in HK104. The second sheet shows the result of viral infection experiments assayed by FISH on various edited strains. The third sheet shows experiments using the *C. elegans rde-11* and *rsd-2* mutants.

**Table S8. LEBV infection of Recombinant Inbred Lines between JU1498 and HK104.** The table indicates the direction of the initial cross for each recombinant inbred line and the phenotyping data. Each line was assayed in three replicates in a given experimental block, for a total of six blocks for the 79 RILs. In each block, the two parents JU1498 and HK104 were infected and scored in parallel, as well as a pre-infected JU1498 culture as additional positive control.

**Table S9. Frequency of the HK104 allele in the sensitive and resistant pools after the JU1498 x HK104 cross.** The chromosomal positions correspond to the *C. briggsae* AF16 Cb4 reference genome.

**Table S10. Genotypes and phenotypes of Near Isogenic Lines.** In the two first sheets, “A” denotes the genotype of AF16 (orange), “B” that of HK104 (blue). The chromosomal coordinates are based on the AF16 Cb4 reference genome.

## References

Andersen, E. C., Rockman, M. V. (2022). Natural genetic variation as a tool for discovery in *Caenorhabditis* nematodes. Genetics 220: iyab156.

Ashe, A., Bélicard, T., et al. (2013). A deletion polymorphism in the *Caenorhabditis elegans* RIG-I homolog disables viral RNA dicing and antiviral immunity. Elife 2: e00994.

Aydin, A., Toliat, M. R., Bahring, S., Becker, C., Nurnberg, P. (2006). New universal primers facilitate pyrosequencing. Electrophoresis 27: 394–7.

Baird, S. E., Davidson, C. R., Bohrer, J. C. (2005). The genetics of ray pattern variation in *Caenorhabditis briggsae*. BMC Evol. Biol. 5: 3.

Bakowski, M. A., Desjardins, C. A., et al. (2014). Ubiquitin-mediated response to microsporidia and virus infection in *C. elegans*. PLoS Pathog 10: e1004200.

Barrett, L. G., Kniskern, J. M., Bodenhausen, N., Zhang, W., Bergelson, J. (2009). Continua of specificity and virulence in plant host-pathogen interactions: causes and consequences. New Phytol 183: 513–29.

Bernstein, M. R., Zdraljevic, S., Andersen, E. C., Rockman, M. V. (2019). Tightly linked antagonistic-effect loci underlie polygenic phenotypic variation in *C. elegans*. Evol Lett 3: 462–473.

Broman, K., Wu, H., Sen, S., Churchill, G. (2003). R/qtl: QTL mapping in experimental crosses. Bioinformatics 19: 889–890.

Casorla-Perez, L. A., Guennoun, R., et al. (2022). Orsay Virus infection of *Caenorhabditis elegans* is modulated by zinc and dependent on lipids. J Virol 96: e0121122.

Castiglioni, V. G., Elena, S. F. (2023). Orsay virus infection increases *Caenorhabditis elegans* resistance to heat shock. bioRxiv https://www.biorxiv.org/content/10.1101/2023.11.02.565305v1.

Chen, K., Franz, C. J., Jiang, H., Jiang, Y., Wang, D. (2017). An evolutionarily conserved transcriptional response to viral infection in *Caenorhabditis* nematodes. BMC Genomics 18: 303.

Chen, S., Zhou, Y., Chen, Y., Gu, J. (2018). fastp: an ultra-fast all-in-one FASTQ preprocessor. Bioinformatics 34: i884–i890.

Coffman, S. R., Lu, J., et al. (2017). *Caenorhabditis elegans* RIG-I homolog mediates antiviral RNA interference downstream of Dicer-dependent biogenesis of viral small interfering RNAs. MBio 8: e00264–17.

Cook, D. E., Zdraljevic, S., Roberts, J. P., Andersen, E. C. (2017). CeNDR, the *Caenorhabditis elegans* natural diversity resource. Nucleic Acids Res 45: D650–D657.

Crombie, T. A., McKeown, R., et al. (2023). CaeNDR, the Caenorhabditis Natural Diversity Resource. Nucleic Acids Res.

Cubillas, C., Sandoval Del Prado, L. E., et al. (2023). The *alg-1* gene Is necessary for Orsay Virus replication in *Caenorhabditis elegans*. J Virol 97: e0006523.

Culp, E., Richman, C., Sharanya, D., Gupta, B. P. (2015). Genome editing in *Caenorhabditis briggsae* using the CRISPR/Cas9 system. Biol Methods Protoc. 5: bpaa003.

Cutter, A. D., Félix, M.-A., Barrière, A., Charlesworth, D. (2006). Patterns of nucleotide polymorphism distinguish temperate and tropical wild isolates of *Caenorhabditis briggsae*. Genetics 173: 2021–2031.

Cutter, A. D., Yan, W., Tsvetkov, N., Sunil, S., Felix, M. A. (2010). Molecular population genetics and phenotypic sensitivity to ethanol for a globally diverse sample of the nematode *Caenorhabditis briggsae*. Mol Ecol 19: 798–809.

Davis, P., Zarowiecki, M., et al. (2022). WormBase in 2022-data, processes, and tools for analyzing *Caenorhabditis elegans*. Genetics 220: iyac003.

Félix, M.-A., Ashe, A., et al. (2011). Natural and experimental infection of *Caenorhabditis* nematodes by novel viruses related to nodaviruses. PLoS Biol. 9: e1000586.

Félix, M. A., Duveau, F. (2012). Population dynamics and habitat sharing of natural populations of *Caenorhabditis elegans* and *C. briggsae*. BMC Biol 10: 59.

Félix, M. A., Wang, D. (2019). Natural viruses of *Caenorhabditis* nematodes. Annu Rev Genet 53: 313–326.

Fodor, A., Riddle, D. L., Nelson, F. K., Golden, J. W. (1983). Comparison of a new wild-type *Caenorhabditis briggsae* with laboratory strains of *Caenorhabditis briggsae* and *Caenorhabditis elegans*. Nematologica 29: 203–217.

Franz, C. J., Renshaw, H., et al. (2014). Orsay, Santeuil and Le Blanc viruses primarily infect intestinal cells in *Caenorhabditis* nematodes. Virology 448: 255–64.

Franz, C. J., Zhao, G., Félix, M.-A., Wang, D. (2012). Complete genome sequence of Le Blanc virus, a third *Caenorhabditis* nematode-infecting virus. J. Virol. 86: 11940.

Frézal, L., Demoinet, E., Braendle, C., Miska, E. A., Félix, M.-A. (2018). Natural genetic variation in a multigenerational phenotype in *C. elegans*. Curr Biol 28: 2588–2596.

Frézal, L., Félix, M. A. (2015). *C. elegans* outside the Petri dish. Elife 4: e05849.

Frézal, L., Jung, H., Tahan, S., Wang, D., Félix, M.-A. (2019). Noda-like RNA viruses infecting *Caenorhabditis* nematodes: sympatry, diversity and reassortment. J. Virol. 93: e01170–19.

Frézal, L., Saglio, M., et al. (2023). Genome-wide association and environmental suppression of the mortal germline phenotype of wild *C. elegans*. EMBO Rep: e58116.

Fusca, D. D., Sharma, E., Weiss, J. G., Claycomb, J. M., Cutter, A. D. (2022). Temperature-dependent small RNA-expression depends on wild genetic backgrounds of *Caenorhabditis briggsae*. Mol Biol Evol 39: msac218.

Guo, X., Zhang, R., Wang, J., Ding, S. W., Lu, R. (2013). Homologous RIG-I-like helicase proteins direct RNAi-mediated antiviral immunity in *C. elegans* by distinct mechanisms. Proc Natl Acad Sci U S A 110: 16085–90.

Guo, X., Zhang, R., Wang, J., Lu, R. (2013). Antiviral RNA silencing initiated in the absence of RDE-4, a double-stranded RNA binding protein, in *Caenorhabditis elegans*. J Virol 87: 10721–9.

Hillier, L. W., Miller, R. D., et al. (2007). Comparison of *C. elegans* and *C. briggsae* genome sequences reveals extensive conservation of chromosome organization and synteny. Plos Biol. 5: e167.

Huson DH, Bryant D (2006) Application of phylogenetic networks in evolutionary studies. Mol Biol Evol 23: 254–267

Inoue, T., Ailion, M., et al. (2007). Genetic analysis of dauer formation in *Caenorhabditis briggsae*. Genetics 177: 809–18.

Jiang, H., Chen, K., Sandoval, L. E., Leung, C., Wang, D. (2017). An evolutionarily conserved pathway essential for Orsay virus infection of *Caenorhabditis elegans*. MBio 8: e00940–17.

Jiang, H., Franz, C. J., Wang, D. (2014). Engineering recombinant Orsay virus directly in the metazoan host *Caenorhabditis elegans*. J Virol 88: 11774–81.

Koboldt, D. C., Staisch, J., et al. (2010). A toolkit for rapid gene mapping in the nematode *Caenorhabditis briggsae*. BMC Genomics 11: 236.

Lazetic, V., Batachari, L. E., Russell, A. B., Troemel, E. R. (2023). Similarities in the induction of the intracellular pathogen response in *Caenorhabditis elegans* and the type I interferon response in mammals. Bioessays 45: e2300097.

Lazetic, V., Wu, F., et al. (2022). The transcription factor ZIP-1 promotes resistance to intracellular infection in *Caenorhabditis elegans*. Nat Commun 13: 17.

Le Pen, J., Jiang, H., et al. (2018). Terminal uridylyltransferases target RNA viruses as part of the innate immune system. Nat Struct Mol Biol 25: 778–786.

Li, H. (2011). A statistical framework for SNP calling, mutation discovery, association mapping and population genetical parameter estimation from sequencing data. Bioinformatics 27: 2987–93.

Li, H., Durbin, R. (2009). Fast and accurate short read alignment with Burrows-Wheeler transform. Bioinformatics 25: 1754–60.

Long, T., Meng, F., Lu, R. (2018). Transgene-assisted genetic screen identifies *rsd-6* and novel genes as key components of antiviral RNA interference in *Caenorhabditis elegans*. J Virol 92: e00416.

Longdon, B., Brockhurst, M. A., Russell, C. A., Welch, J. J., Jiggins, F. M. (2014). The evolution and genetics of virus host shifts. PLoS Pathog 10: e1004395.

Lu, G., Wang, Q., Gao, G. F. (2015). Bat-to-human: spike features determining ‘host jump’ of coronaviruses SARS-CoV, MERS-CoV, and beyond. Trends Microbiol 23: 468–78.

Mao, K., Breen, P., Ruvkun, G. (2020). Mitochondrial dysfunction induces RNA interference in *C. elegans* through a pathway homologous to the mammalian RIG-I antiviral response. PLoS Biol 18: e3000996.

Moffett, P. (2009). Mechanisms of recognition in dominant *R* gene mediated resistance. Adv Virus Res 75: 1–33.

Noble, L. M., Chelo, I. M., et al. (2017). Polygenicity and epistasis underlie fitness-proximal traits in the *Caenorhabditis elegans* Multiparental Experimental Evolution (CeMEE) panel. Genetics 207: 1663–1685.

Ortiz, E. M. (2019). “vcf2phylip v2.0: convert a VCF matrix into several matrix formats for phylogenetic analysis.” from DOI:10.5281/zenodo.2540861. DOI: DOI:10.5281/zenodo.2540861.

Pepin, K. M., Lass, S., Pulliam, J. R., Read, A. F., Lloyd-Smith, J. O. (2010). Identifying genetic markers of adaptation for surveillance of viral host jumps. Nat Rev Microbiol 8: 802–13.

Poplin, R., Chang, P. C., et al. (2018). A universal SNP and small-indel variant caller using deep neural networks. Nat Biotechnol 36: 983–987.

Raj, A., van den Bogaard, P., Rifkin, S., van Oudenaarden, A., Tyagi, S. (2008). Imaging individual mRNA molecules using multiple singly labeled probes. Nature Methods 5: 877–879.

Reddy, K. C., Dror, T., et al. (2017). An intracellular pathogen response pathway promotes proteostasis in *C. elegans*. Curr Biol 27: 3544–3553.

Ren, X., Li, R., et al. (2018). Genomic basis of recombination suppression in the hybrid between *Caenorhabditis briggsae* and *C. nigoni*. Nucleic Acids Res 46: 1295–1307.

Richaud, A., Zhang, G., Lee, D., Lee, J., Félix, M.-A. (2018). The local co-existence pattern of selfing genotypes in *Caenorhabditis elegans* natural metapopulations. Genetics 208: 807–821.

Ross, J. A., Koboldt, D. C., et al. (2011). *Caenorhabditis briggsae* recombinant inbred line genotypes reveal inter-strain incompatibility and the evolution of recombination. PLoS Genet 7: e1002174.

Rothenburg, S., Brennan, G. (2020). Species-specific host-virus interactions: Implications for viral host range and virulence. Trends Microbiol 28: 46–56.

Sarkies, P., Ashe, A., Le Pen, J., McKie, M. A., Miska, E. A. (2013). Competition between virus-derived and endogenous small RNAs regulates gene expression in *Caenorhabditis elegans*. Genome Res 23: 1258–70.

Shaw, C. L., Kennedy, D. A. (2022). Developing an empirical model for spillover and emergence: Orsay virus host range in *Caenorhabditis*. Proc Biol Sci 289: 20221165.

Shirayama, M., Stanney, W., 3rd, Gu, W., Seth, M., Mello, C. C. (2014). The Vasa Homolog RDE-12 engages target mRNA and multiple argonaute proteins to promote RNAi in *C. elegans*. Curr Biol 24: 845–51.

Sowa, J. N., Jiang, H., et al. (2020). The *Caenorhabditis elegans* RIG-I homolog DRH-1 mediates the intracellular pathogen response upon viral infection. J Virol 94: e01173–19.

Sterken, M. G., van Sluijs, L., et al. (2021). Punctuated loci on chromosome IV determine natural variation in Orsay Virus susceptibility of *Caenorhabditis elegans* strains Bristol N2 and Hawaiian CB4856. J Virol 95: e02430.

Stevens, L., Moya, N. D., et al. (2022). Chromosome-level reference genomes for two strains of *Caenorhabditis briggsae:* an improved platform for comparative genomics. Genome Biol Evol 14: evac042.

Stiernagle, T. (2006, February 11, 2006). “Maintenance of *C. elegans*.” Wormbook, from http://www.wormbook.org/. DOI: doi/10.1895/wormbook.1.101.1.

Tanguy, M., Veron, L., et al. (2017). An alternative STAT signaling pathway acts in viral immunity in *Caenorhabditis elegans*. MBio 8: e00924–17.

Tenthorey, J. L., Emerman, M., Malik, H. S. (2022). Evolutionary landscapes of host-virus arms races. Annu Rev Immunol 40: 271–294.

Thomas, C. G., Wang, W., et al. (2015). Full-genome evolutionary histories of selfing, splitting, and selection in *Caenorhabditis*. Genome Res 25: 667–78.

Thompson, J. N. (2005). The geographic mosaic of coevolution. Chicago, University of Chicago Press.

Woolhouse, M., Scott, F., Hudson, Z., Howey, R., Chase-Topping, M. (2012). Human viruses: discovery and emergence. Philos Trans R Soc Lond B Biol Sci 367: 2864–71.

Zdraljevic, S., Walter-McNeill, L., Marquez, H., Kruglyak, L. (2023). Heritable Cas9-induced nonhomologous recombination in *C. elegans*. MicroPubl Biol 2023 DOI: 10.17912/micropub.biology.000775.

